# Persistent postnatal migration of interneurons into the human entorhinal cortex

**DOI:** 10.1101/2022.03.19.484996

**Authors:** Marcos Assis Nascimento, Sean Biagiotti, Vicente Herranz-Pérez, Raymund Bueno, Chun J. Ye, Taylor Abel, Juan S. Rubio-Moll, Jose Manuel Garcia-Verdugo, Eric J. Huang, Arturo Alvarez-Buylla, Shawn F. Sorrells

## Abstract

The entorhinal cortex (EC) is a highly-interconnected hub for multisensory integration and memory processing^1–3^, containing diverse neuronal subtypes^4,5^ including subpopulations that are uniquely spatially-tuned^6,7^. Although many spatial and memory functions develop in infancy, it is considered that neurogenesis and neuronal migration to the EC occurs prenatally. Here we show that the postnatal human temporal lobe contains a prominent stream with large chains of young migrating neurons and many individual neurons breaking away directed into the EC. The EC stream forms between the second and third trimesters of prenatal development when the lateral ventricle walls in the temporal lobe collapse, displacing the subventricular zone (SVZ) and dividing radial glia. At birth, the EC stream follows a path of radial glial fibers in the site of the collapsed ventricle. Migratory chains persist up to 11 months postnatally; however, many individually migrating young neurons can still be detected in the EC at 2 years of age and a few isolated cells at 3 years of age. Within the EC at birth, immature neurons are a mixed population expressing markers of the medial ganglionic eminence (MGE) and caudal ganglionic eminence (CGE), but postnatally rapidly become primarily CGE-derived. Using single-nuclei RNAseq we identified these lineages and found that the MGE-derived neurons matured at earlier postnatal ages compared to those derived from the CGE. The CGE interneurons arriving and maturing the latest included subtypes expressing calretinin (CR), reelin (RELN), and vasoactive intestinal protein (VIP) many of which settle in layer II of the entorhinal cortex. This study reveals that the human EC is still being constructed during the first years of life revealing the largest known postnatal stream of migratory neurons in humans. The protracted postnatal arrival of a diverse population of interneurons could contribute to plasticity^8,9^ and proper excitation-inhibition balance^10,11^ within these highly connected brain circuits.

## Extensive migration in the infant temporal lobe

To investigate postnatal migratory routes in humans we stained human temporal lobe sections at birth for DCX and PSA-NCAM to label migratory neurons. We found an extensive population of large and small clusters of DCX^+^PSA-NCAM^+^ cells surrounding the temporal lobe lateral ventricle (tLV) (**Fig. 1a**) forming a prominent stream that extended up to 1.39 cm in medial-lateral length in coronal sections. This stream extended medio-ventrally below the hippocampus towards the EC and parahippocampal gyrus (PHG) and laterally into the white matter above the inferior temporal gyrus (ITG) and towards the medial temporal gyrus (MTG) (**Extended Data Fig. 1a–c**). Since DCX can be expressed in more mature neurons^4,12,13^, we characterized these putative migrating DCX cells in more detail. At birth, cells in these dense clusters had ultrastructural features consistent with migratory neurons (**Fig. 1b, Extended Data Fig. 1d**): small elongated cell body with leading and trailing processes, scarce cytosol, and an elongated nucleus with compacted chromatin, indicating that these periventricular clusters of DCX^+^PSA-NCAM^+^ cells are chains of migrating young neurons. To identify the extent of this stream along the rostral-caudal level, we mapped its location in coronal sections from the anterior tip of the amygdala to the mid hippocampus (across ∼2 cm). The lateral extension of the stream was only present at anterior levels (**Fig. 1a,c, Extended Data Fig. 1a**). In contrast, the medial extension was present in all anterior-posterior levels examined (**Fig.1c,d, Extended Data Fig. 1a**). At birth, in every level examined, the tLV was surrounded by a network of DCX^+^PSA-NCAM^+^ streams and individual cells that break away from the ventricular-subventricular zone (V-SVZ). We refer to this expansive medially-oriented network of migratory chains and individual cells extending towards the entorhinal cortex as the EC stream.

**Figure 1:**
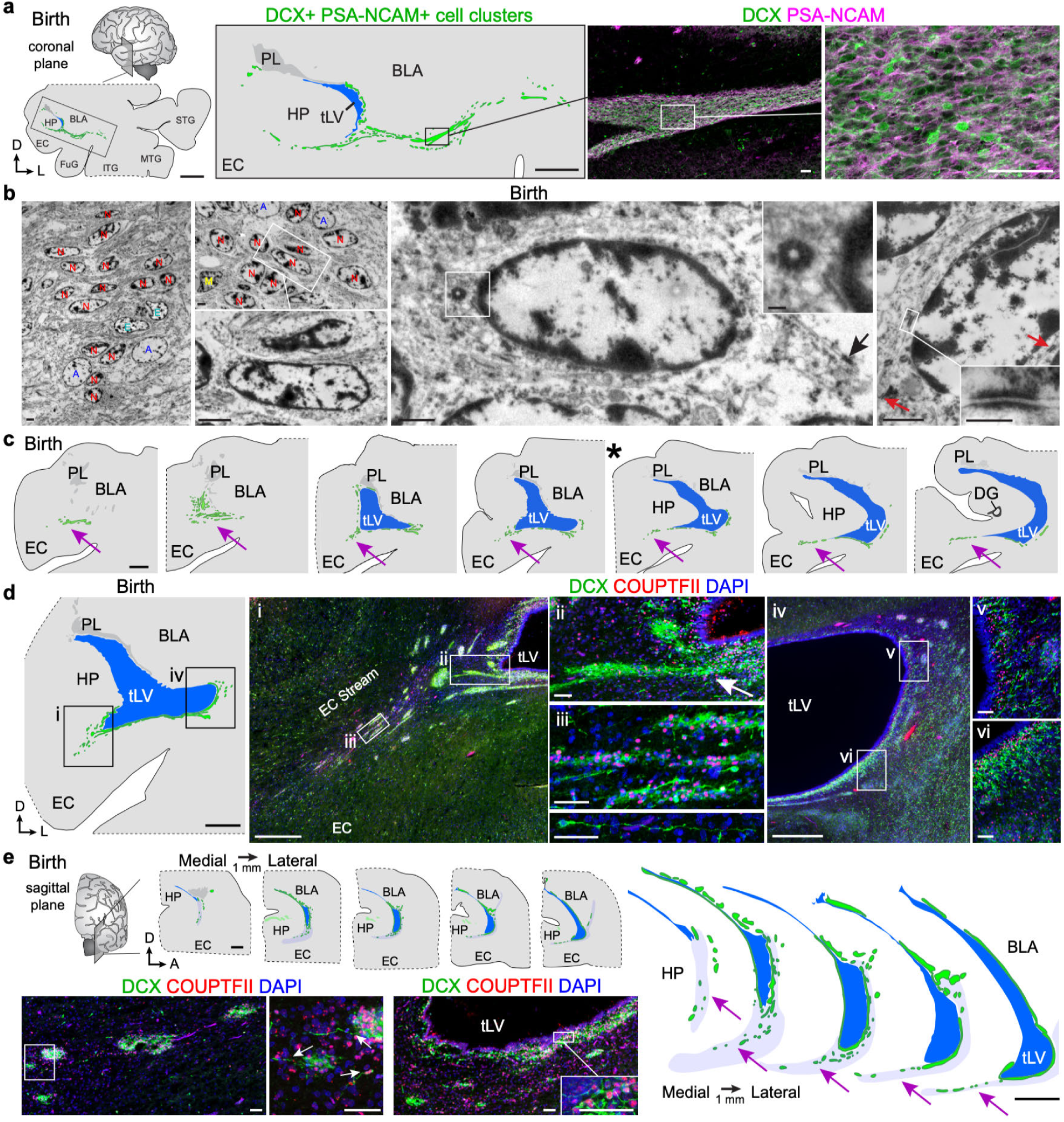
A prominent stream of migratory neurons in the human temporal lobe at birth. a, Coronal map of the human temporal lobe at birth showing the location and immunostaining of dense DCX^+^PSA-NCAM^+^ cell clusters between the EC and BLA. b, Transmission electron microscopy of immature neurons in the human EC stream at birth showing densely packed neurons (red, N) with compacted chromatin and fusiform morphology, surrounded by ependymal cells (cyan, E), astrocytes (blue, A) and microglia (yellow, M). Immature neurons had multiple features of migratory neurons: a centrosome (inset) opposite a trailing process with an adhesion point (arrow). Far right, neuron with multiple adhesion points (inset). c, Serial coronal maps of the medial temporal lobe at birth spaced by 1 mm from the anterior tip of the BLA to the anterior tip of the dentate gyrus showing the location of the EC stream at each level (purple arrows). Asterisk (*) indicates sections adjacent to the one shown in (d). d, Higher magnification map of DCX^+^ COUPTFII^+^ cell clusters at the level indicated by * in (c) with example immunostains of DCX^+^ cells in the EC stream (i-iii) and along the lateral wall of the temporal lobe lateral ventricle (iv-vi). The stream directed to the EC breaks away from the ventricle at its closest point to the EC (arrow). e, Sagittal maps of the medial temporal lobe at birth spaced by 1 mm from the medial edge of the BLA showing the location of dense DCX^+^COUPTFII^+^ cell clusters between the ventricle and EC and immunofluorescence of these cells revealing processes and individual cells extending from the clusters (arrows). Scale bars: 5 mm (a left map), 2 mm (a map inset, c, d, and e maps), 500µm (d i and iv), 50 µm (a right, d ii,iii,v and vi, e bottom), 5 µm (b left and middle left), 1 µm (b middle and right), 200 nm (b middle and right insets).

To identify possible sources for the EC stream, we next asked whether it is labeled by transcription factors associated with different germinal zones. One of the closest germinal zones is the caudal ganglionic eminence (CGE), a massive subpallial germinal zone in humans spanning from the dorsal and ventral caudal forebrain into the temporal lobe. In humans compared to rodents, the CGE contributes proportionally more cortical interneurons than the medial ganglionic eminence (MGE)^14^ and may contribute to increased interneuron diversity in the human cortex^15–19^. At both 18 and 22 gestational weeks (GW), the temporal lobe CGE is present along the dorsolateral tLV and is densely packed with proliferating precursors and young neurons (**Extended Data Figs. 2a–d,g, 3a,c**). This part of the CGE expresses specific combinations of transcription factors, including COUPTFII, SP8, and PROX1 (**Extended Data Figs. 2e,f, 3b**). On the opposing ventromedial wall of the tLV facing the EC, many fewer Ki-67^+^ cells were present (**Extended Data Figs. 2h, 3a**) and the region was filled with DCX^+^DLX2^+^COUPTFII^+^ migratory interneurons (**Extended Data Figs. 2h–j)**. At birth, we mapped the location of Ki-67^+^ cells across a series of temporal lobe coronal sections and identified a hot-spot of proliferating cells in the region where the CGE was prominent at GW22 (**Extended Data Fig. 4a,e**). In this residual CGE we observed clusters of Ki-67^+^COUPTFII^+^ (progenitor) cells closely associated with DCX^+^COUPTFII^+^ cells (immature neurons). Both of these cell types were found more sparsely distributed between the CGE and the EC stream, following the lateral wall of the temporal lobe ventricle and extending into the EC migratory stream (29.0% (324/1119) of CGE Ki-67^+^ cells and 31.3% (50/160) EC stream Ki-67^+^ cells were COUPTFII^+^ **Extended Data Fig. 4b–d**). Given the location of the EC stream along the ventral wall of the tLV, we hypothesized that some, or all, of its cells could be originating in the CGE. Consistently, the majority (88.3%; 339/384) of the DCX^+^ cells in the EC stream expressed COUPTFII, but very few expressed SP8 (2%; 19/967) (**Fig. 1d,e, Extended Data Fig. 1b,c**). We mapped the location of the EC stream at birth in sagittal sections and found similar clusters of DCX^+^COUPTFII^+^ neurons ventral to the hippocampus, between the ventricle and the entorhinal cortex (**Fig. 1e**). Many individual DCX^+^ neurons were present protruding from the streams and in the developing white matter next to and within layers of the neonatal EC (**Fig. 1e, Extended Data Fig. 1b,c**).

## Formation of the EC migratory stream

Given the unique size and structural features of the EC stream at birth, we asked when and how this migratory route forms during gestation. We mapped the location of neural progenitor cells labeled with Ki-67 and SOX2 alongside DCX^+^ neurons between 18 GW and birth in anatomically matched sections at the anterior tip (uncus) of the hippocampus (**Extended Data Fig. 5a–d**). At 18 GW and 22 GW, the temporal lobe CGE was very prominent, containing the vast majority of proliferating cells in our sections (**Extended Data Figs. 2c,d,g, 3a**). At 18 GW the EC was a thin continuous field in the ventral temporal lobe adjacent to the ventricle, which at this stage was open (**Fig. 2a, Extended Data Figs. 2a,c,e,j, 5a**). With the expansion of the hippocampus, CGE, and cortex between 18 and 22 GW, the ventricle in the anterior temporal lobe collapses and the HP and EC ventricular walls come together (**Fig. 2b,c, Extended Data Figs. 3a,b, 5b**). Between 22 and 27 GW, the ventricular region facing the EC had completely fused together, coinciding with additional tissue growth of the surrounding regions (**Extended Data Fig. 5c**). At 27 GW and birth, a distinct population of displaced SOX2^+^ cells was detectable at the former ventricle site, between the hippocampus and EC in the site of the EC stream at birth (**Extended Data Fig. 5c–e**). The ventricular collapse resulted in the displacement of vimentin^+^ (VIM) radial glia (RG) from the wall of the ventricle, distributing them across the EC stream anlage (**Fig. 2c–e**) with their radial processes re-directed medially to take a more tangential orientation along the EC stream (**Fig. 2d**). At birth, displaced HOPX^+^ VIM^+^ RG cells were abundant throughout the EC stream and a subset were still dividing (**Fig. 2d**). It is within this region of the collapsed ventricle and redirected radial processes that chains of neurons migrating around the ventricle coalesce into the EC stream (**Fig. 2d,e**).

**Figure 2:**
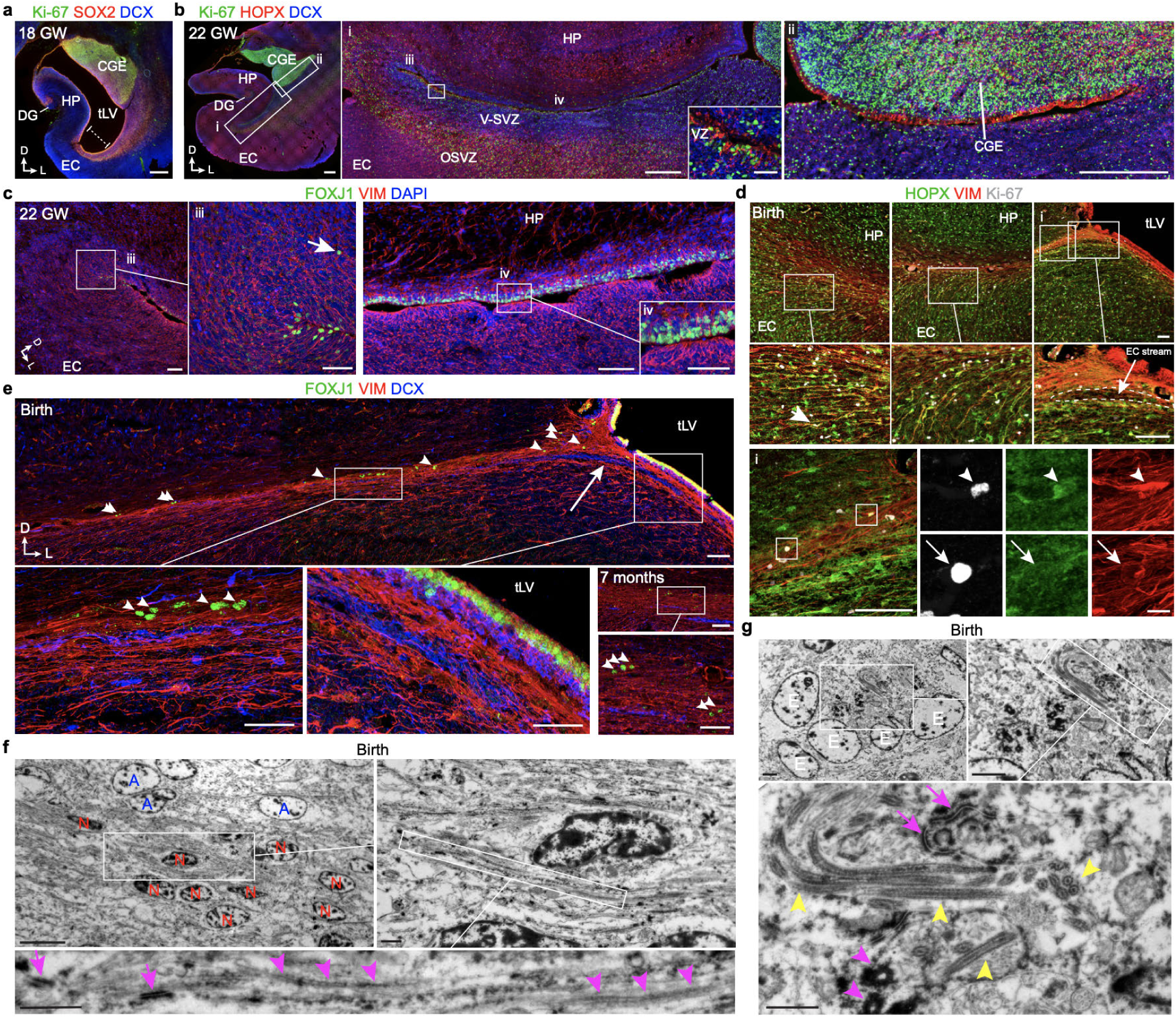
The EC stream migrates along a radial scaffold in a collapsed ventricular zone. **a**, At 18GW, coronal sections show Ki-67^+^SOX2^+^ cells prominent in the CGE and lining the ventricle walls which are open between the medial wall facing the hippocampus and lateral wall facing the cortex (dotted line). Many DCX^+^ cells are found between the CGE and EC. **b**, At 22GW, coronal sections of the same anatomical plane show the medial and lateral VZ pressed closely together. Ki-67^+^ cells are densely clustered in the CGE and present along the lateral wall V-SVZ with DCX^+^ cells filling the V-SVZ between a layer of Ki67^+^HOPX^+^ cells in a temporal lobe outer subventricular zone. **c**, At 22GW, in an immediately adjacent section to (**b**), the ventral tip of the ventricle (region iii in **b**) contains FOXJ1^+^ cells, some displaced in the V-SVZ (arrow) alongside VIM^+^ cells with radial processes. Dorsally along this confluence (region iv in **b**), FOXJ1^+^ cells are present in the medial wall VZ and the lateral wall VZ contains dense VIM^+^ cells and no FOXJ1+ cells (inset). **d**, At birth, the same region no longer has an open ventricle and instead is composed of VIM^+^ radial glial fibers and VIM^+^HOPX^+^ and VIM^+^HOPX^−^ cells, a subset of which are Ki-67^+^ (arrows, insets). **e**, At birth, the DCX^+^ cells in the V-SVZ can be seen turning away from the ventricle toward the EC (arrow) into a VIM^+^ glial corridor. At birth and 7 months, FOXJ1^+^ cells line the wall of the ventricle, and are in groups along the EC stream (arrowheads). **f**, Ultrastructural detail of an immature migratory neuron (red, N) in the EC stream at birth surrounded by glial fibers and cell bodies (blue, A). An adhesion point (zonula adherens, magenta arrows) is visible between this cell and surrounding glial fibers (magenta arrowheads). **g**, A group of 5 multiciliated (yellow arrowheads) ependymal cells (white, E) in the EC stream containing ciliary basal bodies (magenta arrowheads) and cell-cell junctions typical of ependymal cells (magenta arrows). Scale bars: 1 mm (**a, b** left overview), 500 µm (**b** i,ii overviews), 100 µm (**c** iii,iv overviews, **d** top and bottom left, **e** birth and 7 months top), 50 µm (**a** i inset, **c** iii, iv insets, **e** birth insets), 10 µm (**d** i insets, **f** top left), 2 µm (**g** top panels), 1 µm (**f** top right, bottom insets, **g** bottom inset).

To determine whether migratory neurons follow RG fibers, we stained the EC stream at birth and 7 months for VIM, an intermediate filament that is highly expressed in RG. Small chains of DCX^+^ cells were first detected at 27GW in the EC stream (**Extended Data Fig. 5c**), and at birth these clusters were larger and more abundant (**Extended Data Fig. 5d**). The EC stream chains at birth were surrounded by VIM^+^ cell bodies and fibers (**Fig. 2d,e**), resembling the glial tube of the rostral migratory stream^20^(RMS). A similar glial tube surrounding each stream could be seen guiding each tributary of the EC stream away from the wall of the tLV (**Fig. 2d,e, arrows**). Ultrastructural details of migratory neurons in the EC stream at birth confirmed these observations: radial glial fibers and developing astrocytes were observed surrounding chains of elongated migratory neurons that were oriented in the direction of the fibers (**Fig. 2f**).

To further investigate whether the EC stream forms in the location of a collapsed ventricle, we looked for the presence of cells expressing FOXJ1, a transcription factor expressed in multiciliated ependymal cells lining the walls of the ventricle (**Fig. 2c,e**). At 22 GW, FOXJ1^+^ cells were present on the medial wall but not the lateral wall, which contained VIM+ RG fibers, along the compressed ventricular regions (**Fig. 2c**). By birth and at 7 months (when the ventricles have fully collapsed), single, pairs, or small clusters of FOXJ1^+^ cells were observed all along the EC stream (**Fig. 2e**). In this same region we found clusters of multiciliated cells with ultrastructural features of ependymal cells (**Fig. 1b, 2g, Extended Data Fig. 1d**). We conclude that the EC stream forms as an extension from the ventral CGE as chains of migrating young neurons coalesce into a stream that develops between 22 and 27 GW next to the collapsing walls of the tLV.

## The EC migratory stream is depleted by one year of age, and individually migrating neurons persist beyond 2 years in the EC

We next investigated how long the EC stream persists by staining sections of the temporal lobe between birth and 2 years of age for DCX and PSA-NCAM. Dense clusters of DCX^+^PSA-NCAM^+^ cells were present along the temporal lateral ventricle and in the EC stream up to 11 months of age, but were depleted at 14 and 24 months (**Fig. 3a**). At all postnatal ages between birth and 11 months, we observed individual DCX^+^ cells with migratory morphology near the streams, distributed throughout the cortex, and with processes escaping the stream clusters (**Fig. 1d,e, Extended Data Fig. 1b,c**). To determine how long individually migratory DCX^+^ cells persist, we mapped all individual DCX^+^ cells in the medial temporal lobe at birth, and found widespread distribution and a large subset with migratory morphology (**Fig. 3b, Extended Data Fig. 6a–e**). We next stained samples from birth to 3 years of age and drew vector maps of the orientation of the leading process for the subset of individual DCX^+^ cells that displayed clear migratory morphology (a leading process and elongated nuclei) (**Fig. 3b, Extended Data Fig. 6b**). We classified these neurons as oriented either tangentially (parallel) or radially (perpendicular) to the cortical surface (**Fig. 3b**,**c**). At birth, individually migrating cells can be found migrating in both orientations, broadly distributed across the temporal lobe. With increasing age, these cells become more localized to the superficial cortical layers. From birth to 2 years of age, we observed at least twice as many DCX^+^ cells migrating radially as cells migrating tangentially in the EC (66.9% of all migrating neurons at birth; 87.5% at 7 months; 79.9% at 2 years), whereas in the subcortical region, including the site of the EC stream, cells were generally oriented tangentially (50.1% at birth; 58.2% at 7 months; 58.5% at 2 years) (**Fig. 3c**). We also assessed the most frequent orientations of leading processes in sectors of the temporal lobe in sagittal and coronal sectioning planes. This analysis revealed highly polarized migratory neurons oriented toward the cortical surface compared to those closer to the ventricle (**Extended Data Fig. 6c,d**). The number of individually migrating cells sharply decreases in the subcortical regions after birth, disappearing from the EC between 2 and 3 years, consistent with their migratory orientation (**Fig. 3b,c, Extended Data Fig. 6)**. At 3 years of age, the few remaining DCX^+^ cells with a leading process were restricted to sites in the EC and amygdala, where persistent immature neurons are found in humans,^13,21–23^, non-human primates^24–26^, and other mammals^27^ (**Fig. 3b**).

**Figure 3:**
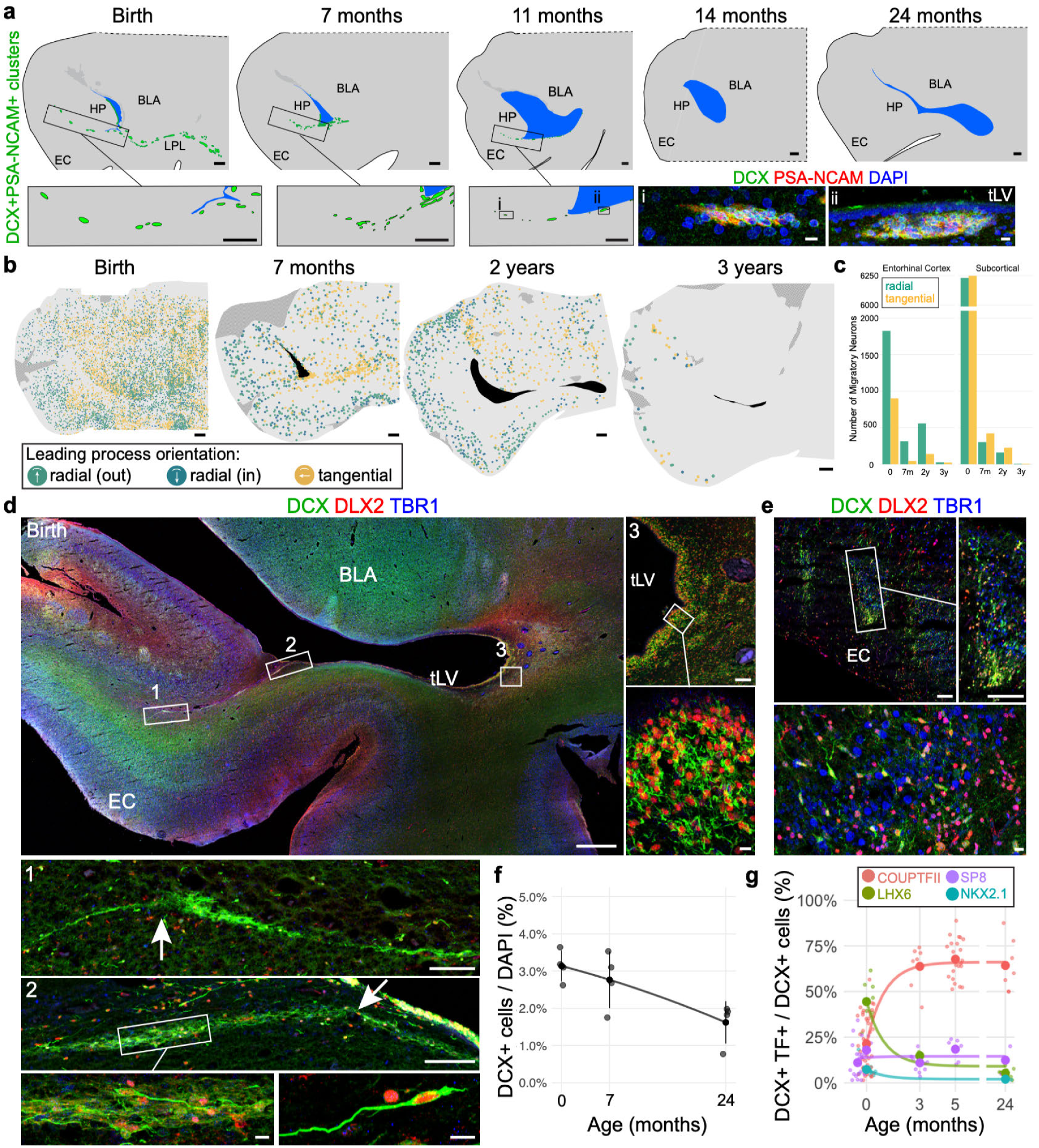
The EC stream supplies CGE-derived interneurons to the EC up to 2 years of age. **a**, Maps of DCX^+^PSA-NCAM^+^ cell clusters in coronal sections of the human medial temporal lobe at the same anatomical level (anterior uncus of hippocampus) from birth to 24 months of age and immunostains showing DCX^+^PSA-NCAM^+^ cell clusters at 11 months of age. **b**, Maps of the orientation of DCX^+^ neurons in coronal sections from birth to 3 years indicating the location of radial (towards or away from the pia) and tangentially-oriented neurons. **c**, Number of neurons with radial and tangential orientations in subcortical white matter and in the EC between birth and 3 years. **d**, Coronal section at birth showing DCX^+^DLX2^+^TBR1^−^ cells in the EC stream (1), the branch-point away from the ventricle (2), and the lateral ventricular wall with a dense field of DCX^+^DLX2^+^TBR1^−^ cells (3). **e**, DCX^+^DLX2^+^TBR1^−^ cells in the EC at birth. **f**, Percentage of total cells expressing DCX at each age. **g**, Percentage of DCX^+^ cells expressing each indicated transcription factor between birth and 2 years of age. Scale bars: 1 mm, (**a** maps, **b, d** top left), 100 µm (**d** 1, 2, and 3 overviews, **e** top panels), 10 µm (**a** bottom right immunostains, **d** bottom panels, 2 inset, 3 inset, **e** bottom panel).

## The EC stream is composed of late-migrating interneurons

Although cells in the EC stream express COUPTFII, this transcription factor is expressed broadly in the amygdala (**Extended Data Fig. 2e,f,j, 3b**), and is insufficient to define them as interneurons^14,22,28,29^. To determine if cells in the EC stream are migrating interneurons, we co-stained DCX^+^ cells for DLX2, a TF required for GABAergic interneuron differentiation^30^. At birth, the majority of DCX^+^ cells in the EC stream (82.4% (1163/1412)) expressed DLX2 (**Fig. 3d, Extended Data Fig. 7a–d**). Between birth and 7 months, following the EC stream back to its connection to the tLV revealed DCX^+^DLX2^+^ cells in chains along the ventricle wall and in the lateral edge of the tLV (**Fig. 3d, Extended Data Fig. 7c,d**). In contrast, at birth we found very few (1.9% (29/1553)) DCX^+^ cells in the EC stream or along the walls of the tLV that expressed the cortical excitatory neuron transcription factor T brain 1 (TBR1)^31^ (**Fig. 3d**). At many levels, we could identify individual DCX^+^ cells breaking away from the stream, oriented toward the EC. Within the EC at birth, 85.5% (868/1015) of DCX^+^ cells expressed DLX2, a similar percentage to the EC stream. In the EC, large collections of DCX^+^DLX2^+^TBR1^−^ cells were observed at the presumptive terminus of the EC stream at birth and at 7 months (**Fig. 3e, Extended Data Fig. 7d**).

## EC DCX^+^ cell identity transitions from a mix of MGE-CGE- to CGE-derived during infancy

Based on the high percentage of EC stream cells expressing DLX2 but not TBR1, we asked if the individual DCX^+^ cells in the EC have the same identity as the cells we observe in the EC stream (**Fig. 3f,g**). At birth we could observe DCX^+^ cells within the EC expressing TBR1^+^; however, these cells did not display migratory morphology, and instead were multipolar (**Extended Data Fig. 7e**). A subset of small DCX^+^TBR1^+^ cells were still present in EC Layer II at much older ages (6 and 13 years, **Extended Data Fig. 7f**). To compare the identity of individual DCX^+^ migratory neurons in the EC with those we observed near the ventricle and within the EC stream, we stained these cells for markers of CGE (COUPTFII, SP8) and MGE (NKX2.1, LHX6). Interestingly, unlike the EC stream which is overwhelmingly COUPTFII^+^, within the EC at birth, 44.4% of the DCX^+^ cells were LHX6^+^ and 21.4% were COUPTFII^+^ (**Fig. 3g**). This suggests that LHX6^+^ MGE-derived interneurons use a different route for their migration into the EC, or arrived earlier in development. With increasing postnatal age, we observed a sharp drop in DCX^+^LHX6^+^ cells and a simultaneous increase in the percentage of DCX^+^COUPTFII^+^ cells in the EC (**Fig. 3g**). The percentage of DCX^+^ cells expressing COUPTFII was steadily ∼65% between 3 months and 2 years of age (**Fig. 3g, Extended Data. Fig. 7g–n**). A smaller population of DCX^+^ cells in the EC expressed SP8, LHX6 or NKX2.1 (12.3%, 5.2%, and 1.9%, respectively at 2 years of age, **Fig. 3g, Extended Data. Fig. 7k–n**). Together, these data indicate that the DCX^+^ cells in the EC stream at birth, and in older ages in the EC, predominantly express the CGE markers DLX2 and COUPTFII.

## Diverse interneuron subtypes migrate into the EC between birth and 3 years

To further characterize the identity and diversity of immature neurons in the EC, we isolated nuclei from snap-frozen samples from the EC at early postnatal ages (33 days to 3 years of age) and performed single-nuclei RNAseq (snRNA-seq). We integrated this developmental dataset with one recently published in the adult EC (50 to 79 years of age)^4^ (**Fig. 4a, Extended Figure 9**). This analysis revealed age-related changes in the transcriptional signature across most cell types in the EC (**Fig. 4b, Extended Data Fig. 9, Supplementary Table 3**).

**Figure 4:**
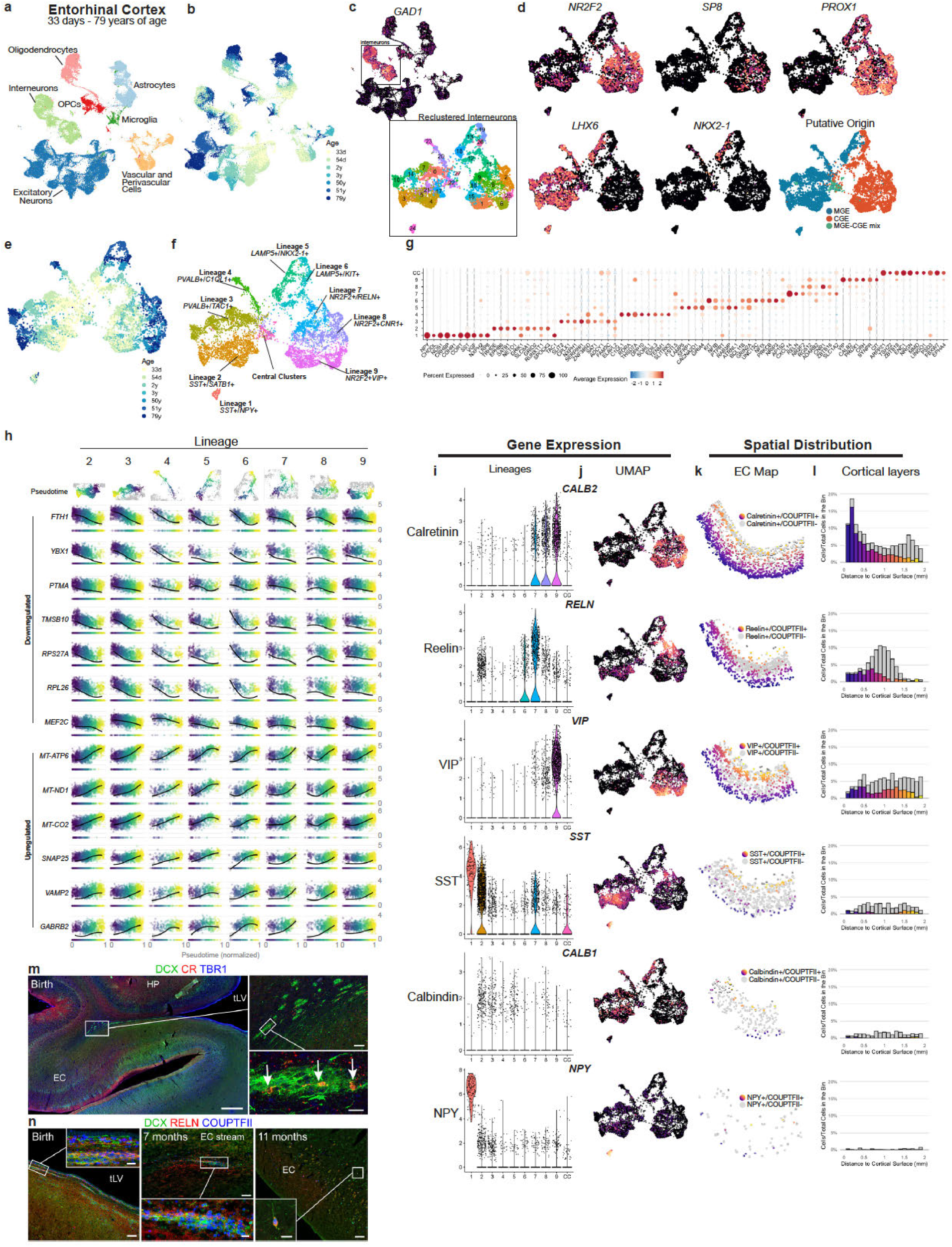
The last neurons arriving in the EC are multiple subtypes of interneurons. **a**,**b**, snRNA-Seq from the EC (33 days to 71 years of age) comprising ∼59,000 cells showing the distribution of main cell types (**a**) and age (**b**). **c**, Reclustering only interneurons reveals multiple subpopulations in the EC. **d**, Expression of the MGE markers *LHX6* and *NKX2-1* and CGE markers *SP8, NR2F2* (COUPTFII), and *PROX1* allows the identification of the putative origin of interneurons. **e**, Dimensionality reduction using UMAP arranges interneurons in a gradient of ages. **f**, Based on the expression of markers of known interneuron subtypes, 9 distinct lineages can be identified. A set of clusters (central clusters) do not correspond to any known interneuron subpopulation in the adult human brain. **g**, Dotplot of the main upregulated DEGs in each lineage and central clusters. **h**, Pseudotime trajectories for each lineage and genes that had the most robust changes along the pseudotime across lineages. **i**,**j**, Violin plots (**i**) and UMAP plots (**j**) showing the expression of classical interneuron markers across the EC interneuron lineages. **k, l**, Spatial distribution of NR2F2^+^ cells expressing the same canonical markers in the EC at birth (**k**) and their frequency distribution across cortical layers (**l**). **m**, CR^+^DCX^+^ cells are present in the EC stream at birth and are TBR1^−^. **n**, RELN^+^DCX^+^ cells are present in the tLV V-SVZ and in the EC stream at birth and 7 months. Scale bars: 1 mm (**m** top left), 100 µm (**m** top right, **n** overview at each age), 20 µm (**m** bottom right, **n** insets).

To better understand interneuron diversity in the EC, we re-clustered them separately and found 28 different clusters (**Fig. 4c)**. These clusters contained cells expressing transcription factors consistent with origins in the MGE (*LHX6* and *NKX2*-*1*) or CGE (*NR2F2, SP8*, and *PROX1*) (**Fig. 4d**). Interestingly, two clusters in the center of the UMAP plot (clusters 22 and 27) had a mixed expression of the markers investigated, suggesting that they consist of a mixed population of MGE and CGE-derived interneurons.

To examine transcriptional changes associated with interneuron maturation, we investigated the distribution of cells coming from different ages across the interneuron clusters. This revealed a continuum across the clusters where inner clusters were from neonate samples 33–54 days old, the middle ring of clusters were from early postnatal samples 2–3 years old, and the outer rim were from adult samples 50-79 years old. (**Fig. 4e**). To identify distinct interneuron subtype lineages within this continuum, we investigated the expression of markers of known interneuron subtypes in the EC and other cortical areas^4,32^, revealing 9 distinct lineages (**Fig. 4f-g**, full list of upregulated differentially expressed genes (DEGs) in **Supplementary Table 4**). The MGE-CGE-mixed clusters in the center of the UMAP plot were not clearly associated with any of the identified maturation trajectories. These central clusters (CC) were composed of cells coming from the earliest ages (33–54 days) and were enriched in genes associated with axon guidance and migration, such as *MEF2C* and *NRP1*. For each lineage, we analyzed possible trajectories using Monocle and used the computed pseudotime estimates to investigate which genes were being differentially expressed along their maturation trajectories. By comparing cells at the early and late stages in their trajectory, we identified genes that have their expression correlating with early and late stages of interneuron maturation. The most robust differences were observed in genes encoding ribosomal proteins (e.g. *RPL26, RPS27A*, **Fig. 4h**; full list in **Supplementary Table 5**), being highly expressed at early stages and downregulated along their maturation trajectory, and ATP-synthesis-associated genes (e.g. *MT-ATP6, MT-CO2, MT-ND1*, **Fig. 4h**; full list in **Supplementary Table 5**), being lowly expressed at early stages and upregulated along their maturation trajectory. The switch from a transcriptional signature associated with protein synthesis to one associated with ATP synthesis was observed in all lineages, irrespective of neuronal type. Genes encoding for the large and small ribosome subunits (RPL and RPS genes) accounted for 54% of all DEGs in interneurons from the youngest age group (33 to 54 days of age, **Extended Data Fig. 9, Supplementary Table 4)**, despite the removal of cells with high ribosomal reads in our QC steps (see Methods). Other genes that had a robust downregulation along all maturation trajectories, were *MEF2C*, associated with neuronal migration and neurite branching. Genes upregulated during interneuron maturation included: *GABRB2, GABRG2, and GABRA1*, GABA receptors, *VAMP2* and *SNAP25*, associated with synaptic vesicle transport and exocytosis (**Fig. 4h, Supplementary Table 5**). Many of the transcriptional changes associated with interneuron maturation happened first in MGE-derived lineages (Lineages 2-5) when compared to CGE-derived lineages (Lineages 6-9), suggesting a late birthdate or delayed maturation CGE-derived neurons in the postnatal EC.

To characterize the anatomical distribution of CGE-derived interneurons arriving in the EC, we mapped the expression of *NR2F2* (COUPTFII). The large majority of DCX^+^ cells in the EC stream at birth were COUPTFII^+^ (88.3%) (**Fig. 3g, Extended Data Fig. 7a**). Consistently, lineages 7, 8, and 9 had a robust expression of *NR2F2*, suggesting that they correspond to the CGE-derived interneurons arriving in the EC. These lineages highly expressed *CALB2, RELN*, and *VIP*, and had low expression of *SST, CALB1* and *NPY*, which are more associated with MGE-derived lineages (**Fig. 4i, j**). Next, we used immunohistochemistry to look at the protein expression of these interneuron subtype markers in the EC at birth. Most neurons expressing the proteins CR, RELN and VIP co-expressed COUPTFII in the upper cortical layers and some RELN^+^/COUPTFII^+^ and VIP^+^/COUPTFII^+^ cells could also be observed in deeper layers (**Fig. 4k, l**). Expressed across lineages 7, 8, and 9, CR was also the marker that was most co-expressed with COUPTFII, and by 2 years of age CR^+^ interneurons comprised 88.1% of the COUPTFII^+^ cells in the EC (**Extended Data Fig. 8b,c**).

To identify young migrating neurons from lineages 7, 8, and 9, we immunostained the EC stream and EC for DCX, CR, and RELN between birth and 2 years. At birth we found DCX^+^CR^+^ cells in the EC stream and in the EC (**Fig. 4m, Extended Data Fig. 8a,b)**. Because CR can be expressed in excitatory neurons, we co-stained for TBR1, and found that the DCX^+^CR^+^ neurons in the EC stream and EC at birth were TBR1^−^ (**Fig. 4m, Extended Data Fig. 8a**). We also found collections of DCX^+^RELN^+^COUPTFII^+^ neurons along the ventral tLV in the V-SVZ at birth, in the EC stream at 7 months, and in the EC at 11 months (**Fig. 4n, Extended Data Fig. 8d–j**). Together these results indicate that the EC stream supplies multiple subtypes of CGE-derived interneurons to the EC including CR^+^ and RELN^+^ interneurons.

## Discussion

Here we show that the temporal lobe in human infants contains a large migratory stream of young neurons directed toward the EC. This stream forms in gestation as the ventricle collapses and is maintained postnatally for at least 11 months, with individual migrating cells continuing to settle between 2–3 years of age, supplying CR^+^, RELN^+^, and possibly other CGE-derived interneurons to the EC.

The scale and persistence of the EC stream may result from continued postnatal progenitor cell proliferation and/or migration from earlier, distant progenitors. At birth in the anatomical location where the CGE is found during embryonic ages (**Extended Data Figs. 2, 3**), we found clusters of Ki-67^+^ cells closely associated with immature neurons, both labeled with COUPTFII, a TF that is highly expressed in the CGE^33,34^ (**Extended Data Fig. 4**). In the embryonic mouse brain, a caudal migration from the CGE into the HP and EC has been described^35^. The migration we describe here in humans could be an expansion in size and duration of a process already present in rodents with the collapsed ventricular zone providing a scaffold. Protracted migration of interneurons has also been described during the first 3 months of life in the human frontal lobes^20,36^. In the case of EC migration, the duration is extensive in postnatal life. We first observed clusters of DCX^+^ cells in the EC stream at 27 GW and more were present by birth and postnatally until 11 months, demonstrating that ∼¼ of the EC stream duration is during gestation, whereas ∼¾ is postnatal. The EC stream is established at the former site of an open ventricle and contains radial glial palisades (**Fig. 2d,e**) that surround the lamina of migratory streams. The V-SVZ is a common migratory route for interneurons, and the rostral migratory stream in rodents emerges in the location of the olfactory ventricle and these cells are surrounded by a glial tube^37^. Similar collapsed ventricular regions have been associated with migratory streams in the dorsolateral ventricles in rabbits^38^. There is recent evidence that pallial progenitors can generate inhibitory neurons that are transcriptionally identical to interneurons of CGE origin^39^, which makes the displaced RG cells also a possible progenitor for these late-arriving neurons.

The EC stream supplies interneurons from the CGE to the EC based on several lines of evidence. The disappearance of DCX^+^PSA-NCAM^+^ and DCX^+^COUPTFII^+^ clusters in the EC stream occurs at younger ages than the disappearance of individual migratory cells in the EC (**Fig. 3a,b**), and the percentage of DCX^+^ cells in the EC expressing COUPTFII increases sharply soon after birth (**Fig. 3g, Extended Data Fig. 7g–n**) suggesting a progression from stream to cortex. We also observe DCX^+^ cells extending their leading process away from the EC stream into the EC (**Figs. 1e, 3d, Extended Data Figs. 1b,c, 6b–d**). Using snRNA-seq, we identified 9 distinct lineages that underwent a continuous change in cell state. These lineages corresponded to maturing MGE and CGE-derived interneurons, with multiple genes associated with their maturation. Maturation trajectories indicate that MGE-derived interneurons mature earlier compared to the lineages derived from the CGE. We observed a switch from a high expression of genes associated with cell migration, cytoskeleton rearrangement, and protein translation to a higher expression of neurotransmitter receptors and ATP synthesis. Interestingly, among the top DEGs in the clusters of the most immature neurons were *ARPP1* or *R3HDM1*; the genes that host miR-128, a key regulator of neuronal migration and branching^40,41^. To identify which EC interneuron subtypes arrive via the EC stream, we examined the lineages using Monocle^42^. This identified CR/VIP^+^ and RELN^+^ cells in layers II and IV, which also expressed the EC-stream marker COUPTFII. Consistently, CR has previously been observed to increase postnatally in the 5th month in the human infant EC^43^ and using immunocytochemistry, we found a subset of DCX^+^ in the EC stream expressing CR **(Fig.)**. RELN, a marker of one of the lineages **(lineage 7, Fig. 4g, i)** was also present in a subset of DCX^+^ cells in the EC stream (**Fig. 4o, Extended Data Fig. 8d–j**). These RELN^+^COUPTFII^+^ cells settle predominantly in layer II of the EC. It is layer II that is initially affected in Alzheimer’s Disease^44,45^, and intrigugingly, RELN^+^ cells in this layer have been observed to accumulate amyloid-beta.

The EC has a critical role in multisensory integration and spatial memory, processes with extensive postnatal development. How the maturation of these processes is associated to the recruitment and maturation of EC-derived interneurons, remains an important question for future research. GABAergic signaling and maturation of local circuit inhibitory cells are critical for periods of enhanced plasticity during development^46–48^. The nature of some of the late arriving interneurons as inhibitory and/or disinhibitory interneuron subtypes may result in a shifting in the excitatory-inhibitory balance^49^ across infancy. The delayed formation and maturation of inhibitory circuits could allow for an extended period of plasticity in the EC and defects in migration or the maturation of these cells may be related to neurodevelopmental disorders affecting this region.

## Acknowledgments

University of Maryland Brain Bank and the NIH Neurobiobank for the help with human samples collection. A.A.B and M.A.N were supported by the National Institute of Neurological Disorders and Stroke (NINDS) (A.A.B and M.A.N: grant R01NS028478, A.A.B: grant P01NS083513). A.A.B was supported by the UCSF Program for Breakthrough Biomedical Research, funded in part by the Sandler Foundation. S.F.S. and S.W.B. were supported by R21MH125367, a Competitive Medical Research Fund award from the University of Pittsburgh Medical Center and start-up funds from the University of Pittsburgh. J.M.G-V. was supported by the Valencian Council for Innovation, Universities, Science and Digital Society (PROMETEO/2019/075) and V.H-P. was supported by the Spanish Ministry of Science, Innovation and Universities (PCI2018-093062).

## Materials and Methods

### Human tissue collection

Thirty-nine post-mortem specimens and one post-operative neurosurgical resection were collected for this study (**Supplementary Table 1**). Tissue was collected with previous patient consent in strict observance of the legal and institutional ethical regulations in accordance with each participating institution: 1. The University of California, San Francisco (UCSF) Committee on Human Research. Protocols were approved by the Human Gamete, Embryo and Stem Cell Research Committee (Institutional Review Board) at UCSF. 2. The Ethical Committee for Biomedical Investigation, Hospital la Fe (2015/0447) and the University of Valencia Ethical Commission for Human Investigation. 3. In accordance with institutional guidelines and study design approval by the committee for research on decedents (CORID) at the University of Pittsburgh. 4. Specimens collected at UPMC had IRB approved research informed consents along with HIPAA authorizations signed by parents or responsible guardians. For infant cases, when the brain is at full term (37 to 40 gestational weeks), we refer to this as “birth”. We collected tissue blocks from the temporal lobe, anteriorly from the amygdaloid complex to the posterior end of the inferior horn of the lateral ventricle. Samples were either flash frozen or fixed in 4% paraformaldehyde (PFA) or 10% formalin for >24h (see **Supplementary Table 1**). Brains were cut into ∼1.5 cm blocks, cryoprotected in a series of 10%, 20%, and 30% sucrose solutions, and then frozen in an embedding medium, OCT. Blocks of the medial temporal lobe were cut into 20 micron sections on a cryostat (Leica CM3050S) and mounted on glass slides for immunohistochemistry.

### Immunohistochemistry

Frozen slides were allowed to equilibrate to room temperature for 3 hours. Some antigens required antigen retrieval (**Supplementary Table 2**), which was conducted at 95°C in 10 mM Na citrate buffer, pH=6.0. Following antigen retrieval, slides were washed with TNT buffer (0.05% TX100 in PBS) for 10 minutes, placed in 1% H2O2 in PBS for 1.5 hours and then blocked with TNB solution (0.1 M Tris-HCl, pH 7.5, 0.15 M NaCl, 0.5% blocking reagent from Akoya Biosciences) for 1 hour. Slides were incubated in primary antibodies overnight at 4°C (**Supplementary Table 2**) and in biotinylated secondary antibodies (Jackson Immunoresearch Laboratories) for 2.5 hours at room temperature. All antibodies were diluted in TNB solution. For most antibodies, the conditions of use were validated by the manufacturer (antibody product sheets). When this information was not provided, we performed control experiments, including no primary antibody (negative) controls and comparison to mouse staining patterns. Sections were then incubated for 30 min in streptavidin-horseradish peroxidase that was diluted (1:200) with TNB. Tyramide signal amplification (PerkinElmer) was used for some antigens. Sections were incubated in tyramide-conjugated fluorophores for 5 minutes at the following dilutions: Fluorescein: 1:100; Cy3: 1:100; Cy5: 1:100. After several PBS rinses, sections were mounted in Fluoromount G (Southern biotech) and coverslipped. Staining was conducted in technical triplicates prior to analysis.

### Fluorescent microscopy, image processing and quantifications

Images were acquired on Leica TCS SP8 or SP5 confocal microscopes using 10x/0.45 NA for tilescans, and 20x/0.75 NA or 63x/1.4 NA objective lenses. Imaging files were analyzed and quantified in Neurolucida software (MBF Bioscience 2017 version). Linear adjustments to image brightness and contrast were made equivalently across all images using Adobe Photoshop (2022). Cells were counted in z-stack images from sections stained for Ki67. Three to five representative images across a minimum of three evenly spaced sections were collected for quantification at each age. Experimental replicates and different co-stains (in addition to the 3-5 sections included for quantifications) were also analyzed for the presence of absence of young neurons or stem cells. Each age has n=1. Counts for cell populations were performed by 3 separate investigators who were blinded to individual cases. For each quantified marker, counts were repeated by different investigators for reproducibility. Fluorescence signal for single reactivity and co-localization of immunoreactivity was counted individually using the markers function in the Neurolucida imaging software. The quantification of data was performed with GraphPad Prism (v9) and R (v4.1).

### Mapping and quantification of young migrating neurons

To generate maps of the location of the somas of different cell types, 2D tilescans were generated using Neurolucida (10x/0.45NA or 20x/0.8NA objective) and contours or markers were placed to indicate the location of each cell type. To generate rose histograms, the angle of the leading process was measured using ImageJ. Leading processes were identified as one single process extending from a DCX+ cell soma. The frequency of DCX+ cell orientations within each segment of the tissue was plotted as a 360-degree histogram using R, and then histograms were overlaid on the map of the tissue within Adobe Illustrator (2022). For vector orientation maps, to determine the angle of migration of each neuron, the angle formed by the shortest path connecting the cell body to the cortical surface and the cell’s leading process was measured. If the resulting angle was between -45° and 45°, cells were classified as migrating radially towards the cortical surface (radial(in)). Cells were classified as migrating tangentially if the angle was between 45° and 135° or -45° and -135°. Cells were classified as radially migrating away from the ventricle if the angle was between -135° and 135° (radial(out)).

### Transmission electron microscopy

Temporal lobe tissue fixed with 2.5% glutaraldehyde-2% PFA in 0.1 M phosphate buffer (PB) was transversely sectioned at 200 µm using a Leica VT1200S vibratome (Leica Microsystems GmbH, Heidelberg, Germany). Slices were further post-fixed in 2% osmium tetroxide in 0.1 M PB for 1.5 hours at room temperature, washed in deionized water, and partially dehydrated in 70% ethanol. Samples were then incubated in 2% uranyl acetate in 70% ethanol in the dark for 2.5 h at 4 ºC. Brain slices were further dehydrated in ethanol followed by propylene oxide and infiltrated overnight in Durcupan ACM epoxy resin (Fluka, Sigma-Aldrich, St. Louis, USA). The following day, fresh resin was added and the samples were cured for 72 h at 70 ºC. Following resin hardening, semithin sections (1.5 µm) were obtained and lightly stained with 1% toluidine blue for light microscopy. Ultrathin sections (70–80 nm) were obtained with a diamond knife using a Ultracut UC7 ultramicrotome (Leica), stained with lead citrate and examined under a FEI Tecnai G2 Spirit transmission electron microscope at 80 kV (FEI Europe, Eindhoven, Netherlands) equipped with a Morada CCD digital camera (Olympus Soft Image Solutions GmbH, Münster, Germany).

### Tissue Processing for snRNA-seq

Snap-frozen samples of entorhinal cortex were obtained from the University of Maryland Brain Bank/NIH Neurobiobank (for Case List see **Supplementary Table 1**). Samples were sectioned (50um) in a cryostat and immediately transferred to -80C for RIN measurement and nuclei extractions. For each sample, total RNA was extracted from 40mg of tissue using TissueRuptor II (Qiagen) and the RNA integrity was measured on the Agilent 2100 Bioanalyzer using the RNA Pico Chip Assay. Only samples with RIN > 7 were used (mean: 7.78).

### Nuclei isolation and snRNA-seq

Frozen sections were transferred from tubes in dry ice to ice cold lysis buffer (0.01M Tris-HCl, 0.14M NaCl, 1mM CaCl2, 0.02M MgCL2, 0.03% Tween-20, 0.01% BSA, 10% Nuclei Ez Lysis Buffer (Sigma), 0.2U/ul Protector RNAse inhibitor (Sigma) in DEPC-treated water) and the tissue was dissociated using a glass dounce homogenizer (Thomas Scientific, Cat # 3431D76). Five samples were processed in parallel and each homogenate was transferred to a separate 30 mL thick wall polycarbonate ultracentrifuge tube (Beckman Coulter, Cat # 355631). Sucrose solution (1.8 M sucrose, 3 mM MgAc2, 1 mM DTT, 10 mM Tris-HCl in DEPC-treated water) was added at the bottom of the tube and homogenates were centrifuged at 107,000 x g for 2.5 hours at 4C. The supernatant was discarded, and the nuclei pellet was incubated in 200 uL of wash/ressuspension buffer (WRB: 1% BSA, 0.2U/ul Protector RNAse inhibitor in DEPC-based PBS) for 20 min on ice before resuspending the pellet. Resuspended nuclei were filtered twice through a 30 um strainer. Each one of the five nuclei suspensions was incubated with its own unique CellPlex Cell Multiplexing Oligo (CMO) barcode (10x Genomics) for 5 minutes followed by a 500 x g centrifugation at 4C for 10 minutes. After centrifugation, supernatant was discarded and nuclei were ressuspended in WRB. After two more centrifugations/WRB washes to remove unbound CMOs, 10ul of each nuclei suspension was stained with DAPI and counted on a hematocytometer. Samples were pooled together and diluted to 2,000 nuclei/ul. A target capture of 30,000 nuclei/well in two wells of a G chip (10x Genomics) was used. Gene expression and barcode libraries were prepared in parallel using the Chromium Next GEM Single Cell 3’ Kit v3.1 (10x Genomics) according to manufacturer’s instructions and sequenced in a Novaseq 6000 system (Illumina). A detailed step-by-step protocol can be found on protocols.io: dx.doi.org/10.17504/protocols.io.kqdg369keg25/v1.

### Read alignment, QC and integration with adult EC dataset

Sequencing reads were aligned using the CellRanger Count v6.1.2 pipeline (10x Genomics) to the human (GRCh38) 2020-A reference. FASTQ files from a published dataset exploring the adult EC^4^ was processed with the same pipeline. The two datasets were integrated using the CellRanger Aggregate v6.1.2 pipeline (10x Genomics) and each dataset was equalized to have the same average read depth (see **Extended Figure 9d**). Cells with either few transcripts (bottom 5%), high percentage of reads belonging to mitochondrial genes (>10%), a high percentage of reads belonging to ribosomal genes (>10%) or were not assigned as singlets by freemuxlet were removed during our QC steps. Reads were normalized to the total reads per nucleus and log transformed.

### Sample demultiplexing and doublet removal

Freemuxlet, a genetic demultiplexing tool in the popscle suite (https://github.com/statgen/popscle), was used to genetically demultiplex cell donors. The bam files generated in the CellRanger Count pipeline were used as inputs for freemuxlet and a customized VCF file from the 1000 genomes data filtered for high variant confidence, Minor Allele Frequency (MAF 0.01), and exonic variants as a reference for SNPs. Freemuxlet assigns each droplet barcode to a single donor, or multiple donors in case a droplet contains cells from distinct samples. Donor to sample matching was done by assessing the amount of each CMO in all cells of a designated donor.

### Dimensionality reduction and clustering

We normalized the data using regularized negative binomial regression (SCTransform) and principal component (PC) analysis was performed on Seurat v3.2.3 ^50^ and 100 PCs (integrated dataset with all cells) or 40 PCs (reclustered data with interneurons only) were used for Uniform Manifold Approximation and Projection (UMAP) dimension reduction and clustering.

### Pseudotime estimation

For pseudotime estimation we used Monocle 3^51^. In short, we subset the cells belonging to each lineage and used Monocle to learn the trajectory graph with the option to allow closed loops in the trajectory disabled. As the neurons coming from the youngest ages were at the center of the UMAP plot (**Fig. 4e, f**), we selected the cells closest to the center of the UMAP plot as the root cells for each lineage. Pseudotime for Lineage 1 and central clusters was not calculated due their small cluster sizes.

### Differential gene expression

To identify differentially expressed genes between clusters and lineages we used a Wilcoxon rank sum test in Seurat v3.2.3 and filtered for genes present in at least 50% of cells in a cluster/lineage/age group, in less than 30% of the cells outside its cluster/lineage/age group, a minimum average fold-change of 0.25 and a maximum false-discovery rate of 5%. For identifying genes differentially expressed along the pseudotime, for each lineage we compared cells with a normalized pseudotime < 0.33 with cells with a normalized pseudotime > 0.66.

## Extended Data Figures

**Extended Data Fig. 1:**
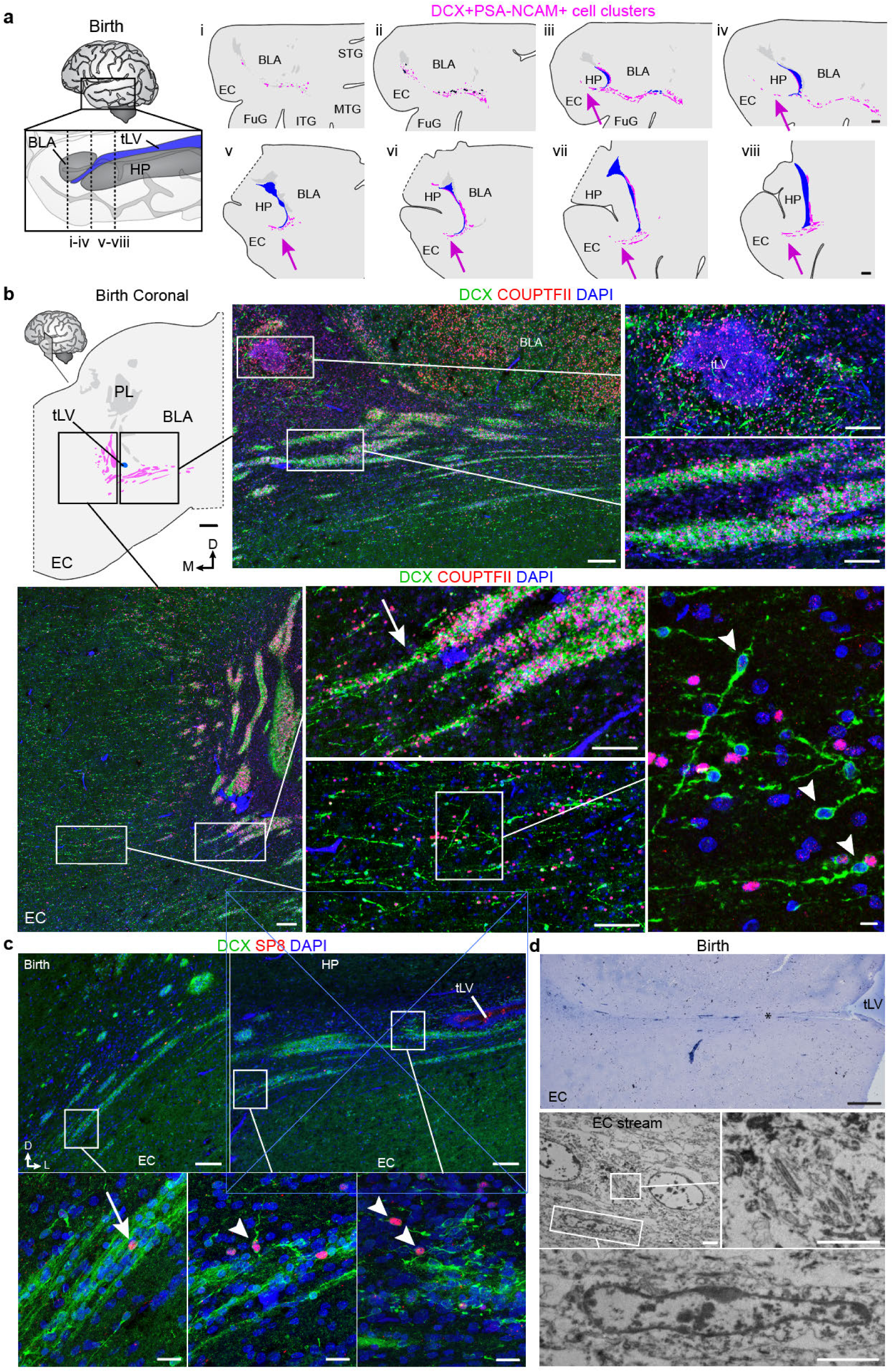
The entorhinal cortex stream of migratory neurons spans the human medial temporal lobe at birth. **a**, Map at birth of DCX^+^PSA-NCAM^+^ cell clusters in a coronal series of sections of the medial temporal lobe. Clusters distributed along a medial-lateral orientation are present ventral to the basolateral amygdala between the EC and medial temporal gyrus (i-iv). The EC stream (arrow) is visible from the first section containing the temporal lobe ventricle and at every section examined posteriorly across the hippocampus (iii-viii). **b**, Map at birth of DCX^+^COUPTFII^+^ cell clusters in a coronal section at the level of the anterior tip of the temporal lobe lateral ventricle (tLV). DCX^+^COUPTFII^+^ cells are primarily found within and emanating from dense streams (arrow) at this age and individual DCX^+^ cells are primarily DCX^+^COUPTFII^−^ (arrowheads). **c**, DCX^+^SP8^+^ cells are rare in the EC stream at birth (arrow), and individual DCX^+^ cells are more frequently SP8^+^ (arrowheads). **d**, Semithin section stained with toluidine blue showing the EC stream at birth (left), and TEM of cells displaying motile cilia (inset) and an elongated neuron with ultrastructural features of a migrating cell in the EC stream at birth (right). Scale bars: 1 mm (**a, b** map), 500 µm (**d** top), 200 µm (**b** top and bottom left overviews), 100 µm (**b** top right insets, bottom middle insets), 20 µm (**c** bottom insets), 10 µm (**b** bottom right), 2 µm (**d** three bottom panels).

**Extended Data Fig. 2:**
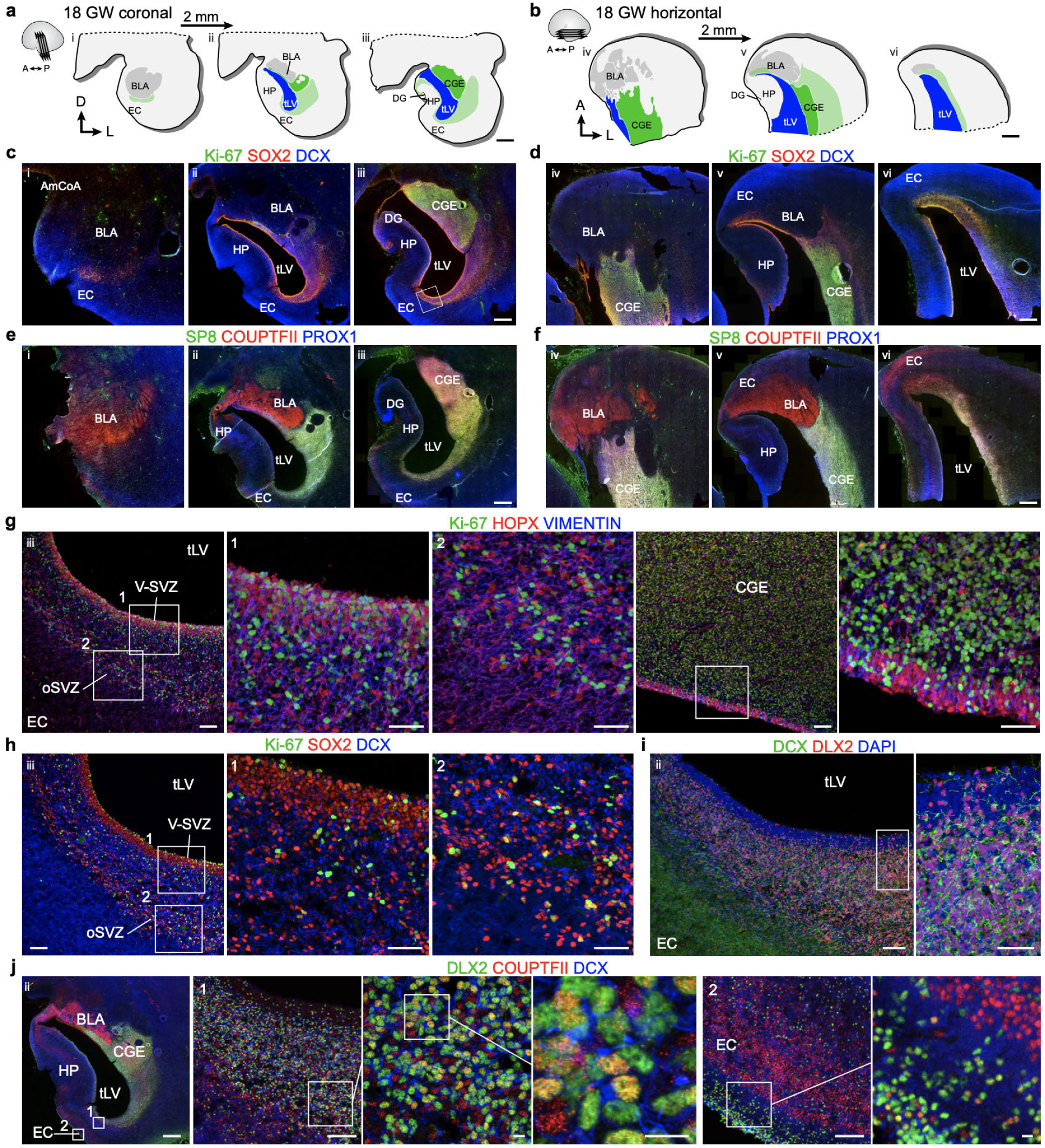
At 18 GW the temporal lobe ventricle is surrounded by dividing progenitors in the CGE and DCX^+^DLX2^+^COUPTFII^+^ neurons extending toward the EC. **a**,**b**, Diagrams of coronal (**a**) and horizontal (**b**) sections of the anterior temporal lobe at 18GW showing the location of the CGE relative to the ventricle and EC. **c**,**d**, Immunostaining for Ki67^+^SOX2^+^ progenitors and DCX^+^ young neurons in sections corresponding to those diagrammed in (**a**,**b**). (Note: level ii corresponds to **Extended Data Fig. 5a** and level iii corresponds to **Fig. 2a**.) **e**,**f**, Immunostaining of sections immediately adjacent to those in (**c**,**d**) for SP8, COUPTFII, and PROX1. **g**, On the ventromedial wall of the temporal lobe ventricle, many Ki67^+^ cells expressing VIM and HOPX are present in the V-SVZ and OSVZ. Similar cells are observed on the opposing wall in the CGE. **h**,**i**, The same region in (**g**) facing the EC contains Ki67^+^SOX2^+^ cells mixed with DCX^+^ neurons (**h**), the vast majority of which are DLX2^+^ (**i**). **j**, Immunostaining of a section adjacent to the middle section in the coronal series (**c–f**) reveals a mixed population of COUPTFII^+^ and COUPTFII^−^ cells expressing DLX2 along the ventricle facing the EC. Scale bars: 2 mm (**a, b**), 1 mm (**c–f, j** left panel), 100 µm (**g** left overview, right CGE overview, **h** left panel, **i** left panel, **j** 1 and 2 left panels), 50 µm (**g** 1, 2, CGE right panel, **h** 1, 2, **i** right panel), 10 µm (**j** 1, two right panels, 2 right panel).

**Extended Data Fig. 3:**
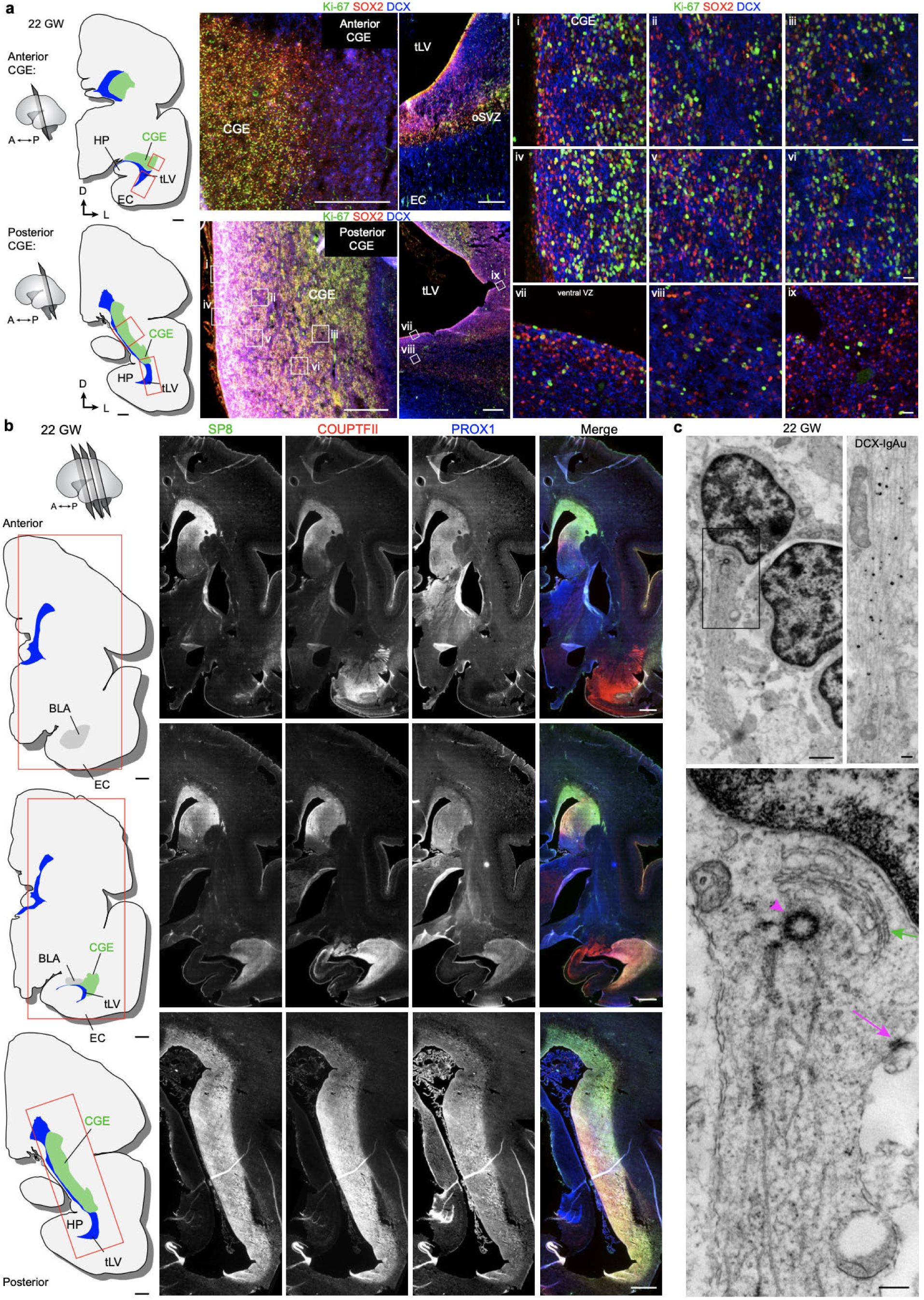
At 22 GW the CGE contains Ki67^+^SOX2^+^ progenitors and many SP8^+^, COUPTFII^+^, and PROX1^+^ cells. **a**, Diagrams of coronal sections of one hemisphere of the human brain at 22 GW at one anterior and one posterior level of the CGE. Red boxes indicate inset locations of immunostaining for Ki67^+^SOX2^+^ cells. Immature DCX^+^ neurons are present in clusters between the Ki67^+^SOX2^+^ cells (i-vi). In the ventral V-SVZ, fewer Ki67^+^SOX2^+^ cells are visible (vii-ix). **b**, Diagrams of coronal sections at three cross-sections across the temporal lobe and immunostains for SP8, COUPTFII, and PROX1 at each level showing that each marker is highly expressed throughout the CGE. **c**, Ultrastructure of a migratory young neuron at 22 GW in the temporal lobe CGE. This cell has a classical localization of the Golgi (green arrow) and centrosome (magenta arrowhead) in the leading process filled with microtubules and displays an adherens junction (magenta arrow). Immunogold labeling for DCX reveals processes filled with microtubules at this age. Scale bars: 2 mm (**a** maps, **b** all panels), 500 µm (**a** left immunostaining overviews), 20 µm (**a** right panels i-ix), 1 µm (**c** top left), 200 nm (**c** top right, bottom panels).

**Extended Data Fig. 4:**
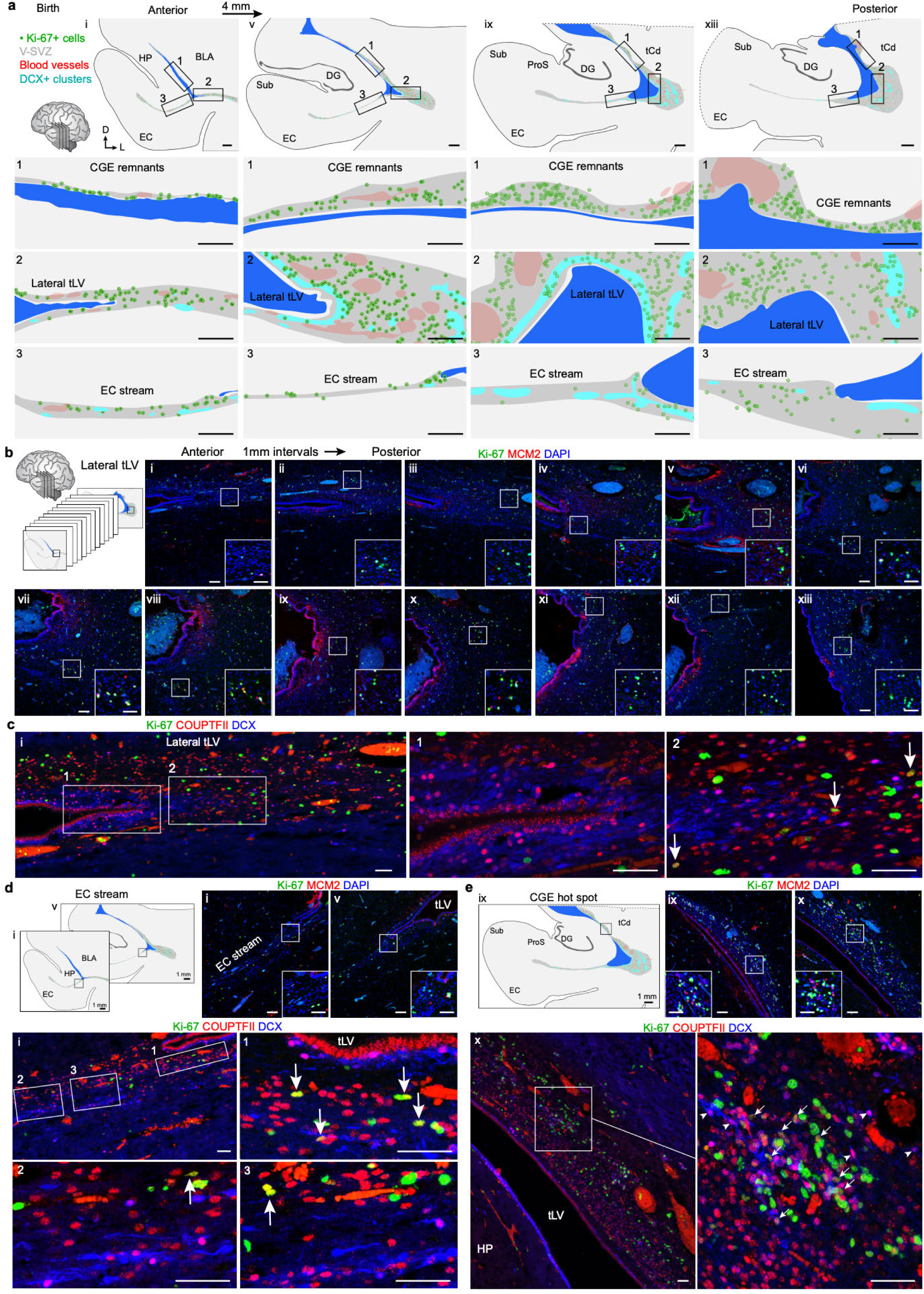
Clusters of dividing progenitors and immature neurons persist along the walls of the temporal lobe at birth. **a**, Maps of coronal sections of the temporal lobe lateral ventricle (tLV) indicating Ki-67^+^ cells (green dots), the V-SVZ region (grey), blood vessels (BV, red), and DCX^+^ cell clusters (cyan). Sections are spaced by 4 mm along the rostral-caudal axis beginning at the anterior/ventral uncus of the hippocampus (i), extending across the rostral dentate gyrus (DG) (v), and more caudally across the DG (ix, xiii). The three inset regions are (1) remnants of the CGE along the dorsolateral wall of the tLV, (2), the most lateral extension of the tLV, and (3) the EC stream. **b**, Dividing Ki-67^+^ cells expressing MCM2 are present along the lateral extension of the tLV in the temporal lobe in the V-SVZ. Sections are spaced by 1 mm. **c**, A stream of DCX^+^ neurons wraps around the lateral extension of the tLV (1) and is present near Ki-67^+^COUPTFII^+^ cells (2) which are a subset of the Ki-67^+^ population in the lateral wall at birth (arrows). **d**, A subset of Ki-67^+^ cells in the EC stream express COUPTFII (arrows) shown at level (i). **e**, A cluster of Ki-67^+^COUPTFII^+^ cells (arrows) and DCX^+^COUPTFII^+^ cells (arrowheads) in the residual CGE at level (x). Abbreviations: BLA basolateral amygdala, CGE caudal ganglionic eminence, DG dentate gyrus, EC entorhinal cortex, HP hippocampus, ProS prosubiculum, Sub subiculum, tCd tail of the caudate nucleus, tLV temporal lobe lateral ventricle. Scale bars: 1 mm (**a** top maps, **d** maps, **e** maps), 500 µm (**a** insets 1,2 and 3), 100 µm (**b** i–xiii overviews, **d** i and v overviews, **e** ix and x overviews), 50 µm (**b** i–xiii insets, **c, d** i, v insets and i 1–3, **e** ix, x insets and x bottom).

**Extended Data Fig. 5:**
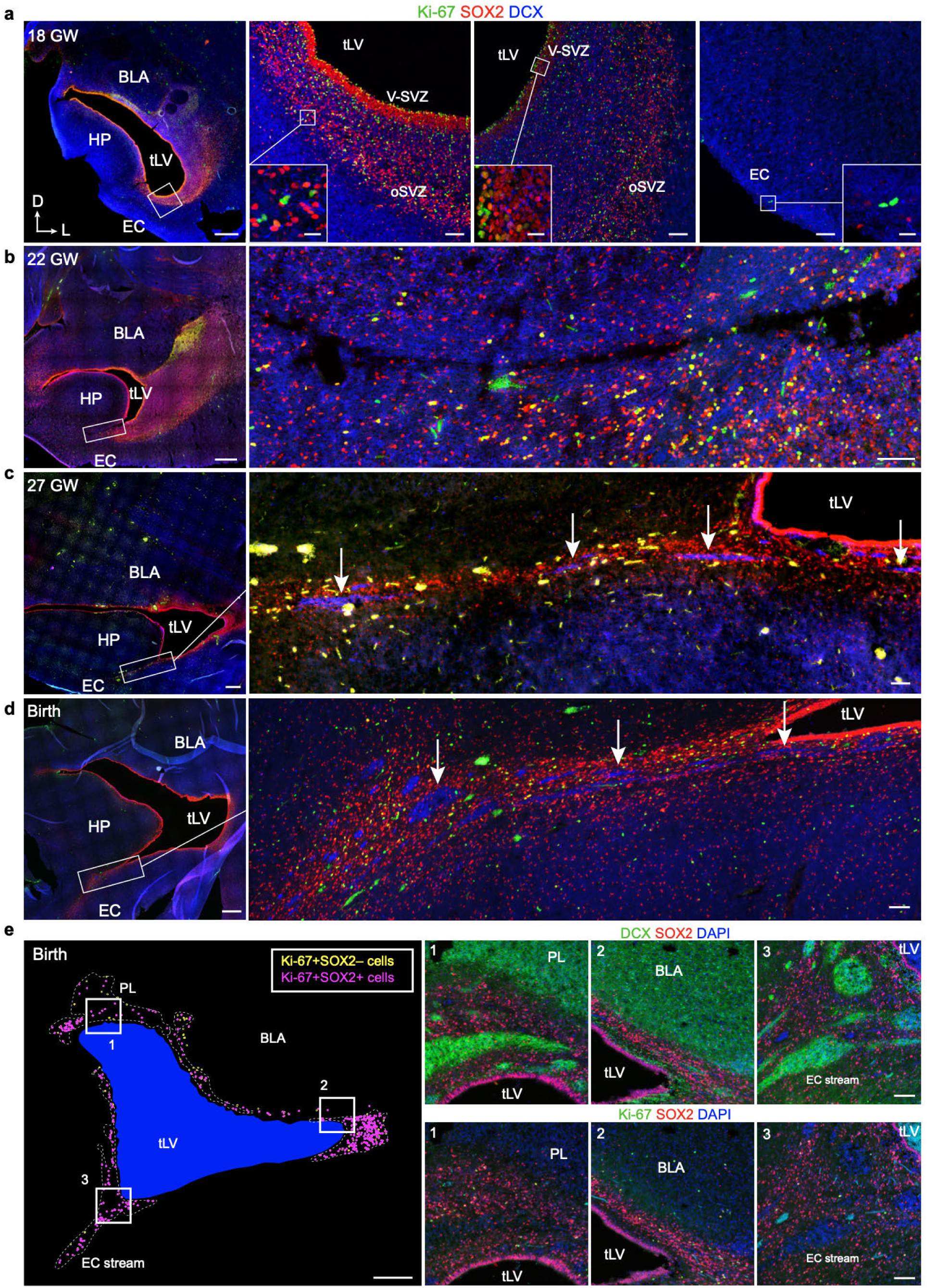
The EC stream forms between 22 and 27 GW. **a–d**, Anatomically matched coronal sections of the medial temporal lobe at the rostral tip of the uncus of the hippocampus across ages. At 18 GW (**a**), Ki-67^+^SOX2^+^ cells line the walls of the temporal lobe ventricle (same sectioning level shown in (ii) in **Extended Data Fig. 2c**), and the medial wall facing the EC and hippocampus has fewer of these cells and no anatomical features separating the hippocampus and EC. At this age, there is a slight tissue protrusion of the hippocampus in the lateral direction towards the ventricle, a feature that becomes rapidly more prominent in the next weeks. At 22 GW (**b**), the hippocampus and surrounding tissue have all grown larger and a seam has begun to form between the hippocampus and EC. At this age, there is no heightened accumulation of Ki-67^+^ SOX2^+^ or DCX^+^ cells within this region. At 27 GW (**c**), this region has formed a higher concentration of Ki67^+^SOX2^+^ cells as well as DCX^+^ cell clusters indicating that the EC stream has formed by 27 GW. At birth (**d**), the density of both the Ki-67^+^ SOX2^+^ cells and the DCX^+^ cell clusters is increased, with many of both populations present within the EC stream extending from the medio-ventral tLV to the EC. **e**, Map of the location of Ki-67^+^SOX2^−^ (yellow) and Ki-67^+^SOX2^+^ (magenta) cells surrounding the tLV at birth. Most Ki-67^+^ cells at this age are SOX2^+^, and are found in the paralaminar nucleus of the amygdala (PL) (1), the lateral V-SVZ (2), and in the EC stream (3). Scale bars: 1 mm (**a–e** all left panel overviews), 100 µm (**a–d** right panel overviews, **e** 1–3), 20 µm (**a** right insets).

**Extended Data Fig. 6:**
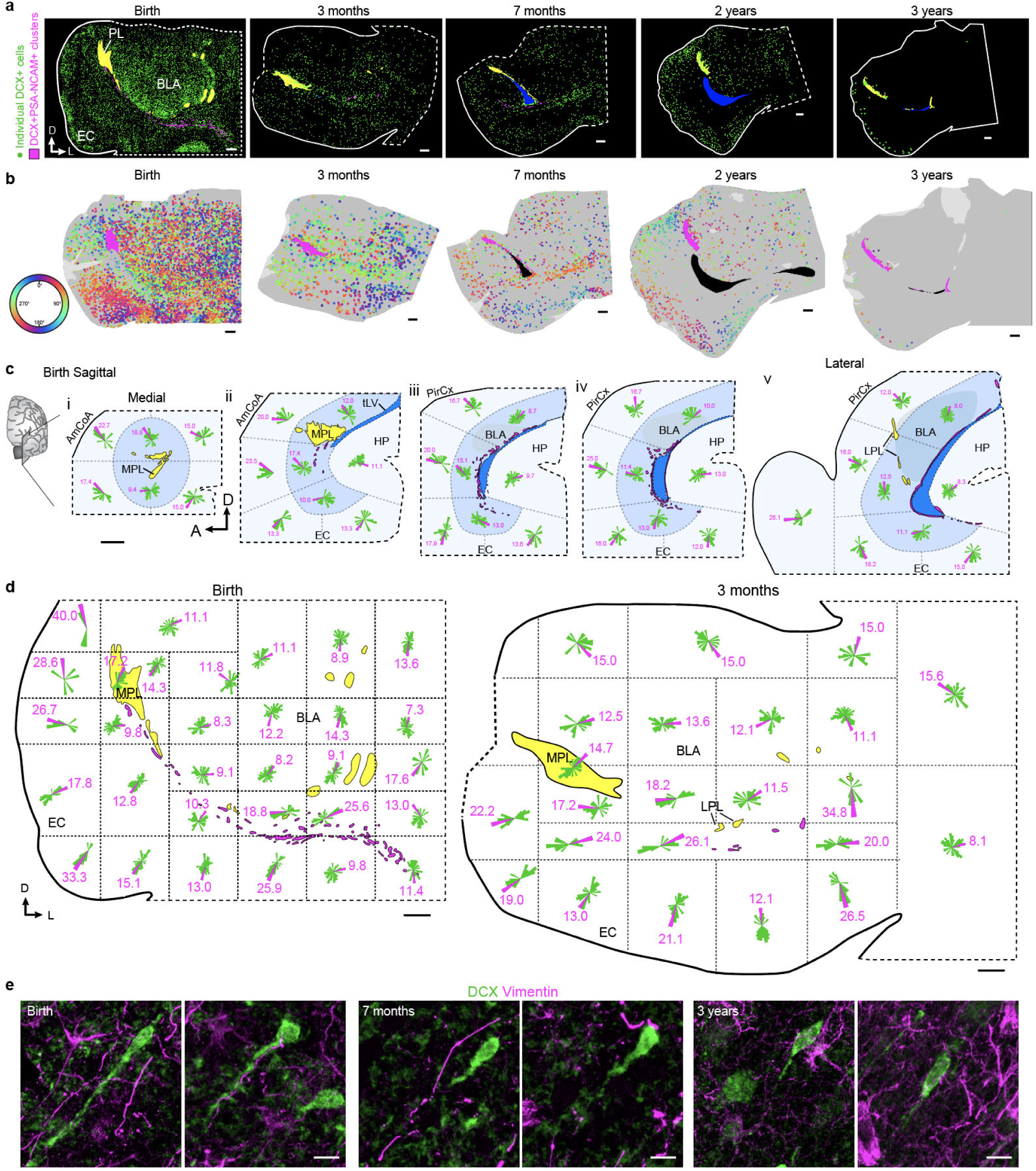
Individual DCX^+^ migratory neurons migrate into the EC from birth to 3 years of age. **a**, Maps of DCX^+^ neurons (green) in coronal sections between birth and 3 years. **b**, DCX^+^ cell orientations between birth and 3 years of age in the same coronal sections indicating radial and tangential orientations in **Fig. 3b**. Here the orientation of the leading process is depicted in 360 degrees by the color of the vector. **c**, Sagittal maps at birth from medial (i) to lateral (v) segmented into sectors (dotted lines). Within each sector, individual DCX^+^ cell orientations are plotted on a 360 degree (rose) frequency histogram. The magenta orientation indicates the percentage of cells that fall into the most frequently occupied orientation within the sector. **d**, Coronal maps at birth and 3 months of age segmented into sectors (dotted lines) containing rose frequency histograms. **e**, Individual DCX^+^ neurons at birth, 7 months, and 3 years of age in the EC and their relationship to VIM^+^ processes. At each age, DCX^+^ cells with migratory features are not associated with VIM^+^ cellular processes. At birth, an example of a DCX^+^ cell with a rounded soma is shown associated with a VIM^+^ process. Scale bars: 2 mm (**c**), 1 mm (**a**,**b**,**d**), 10 µm (**e**).

**Extended Data Fig. 7:**
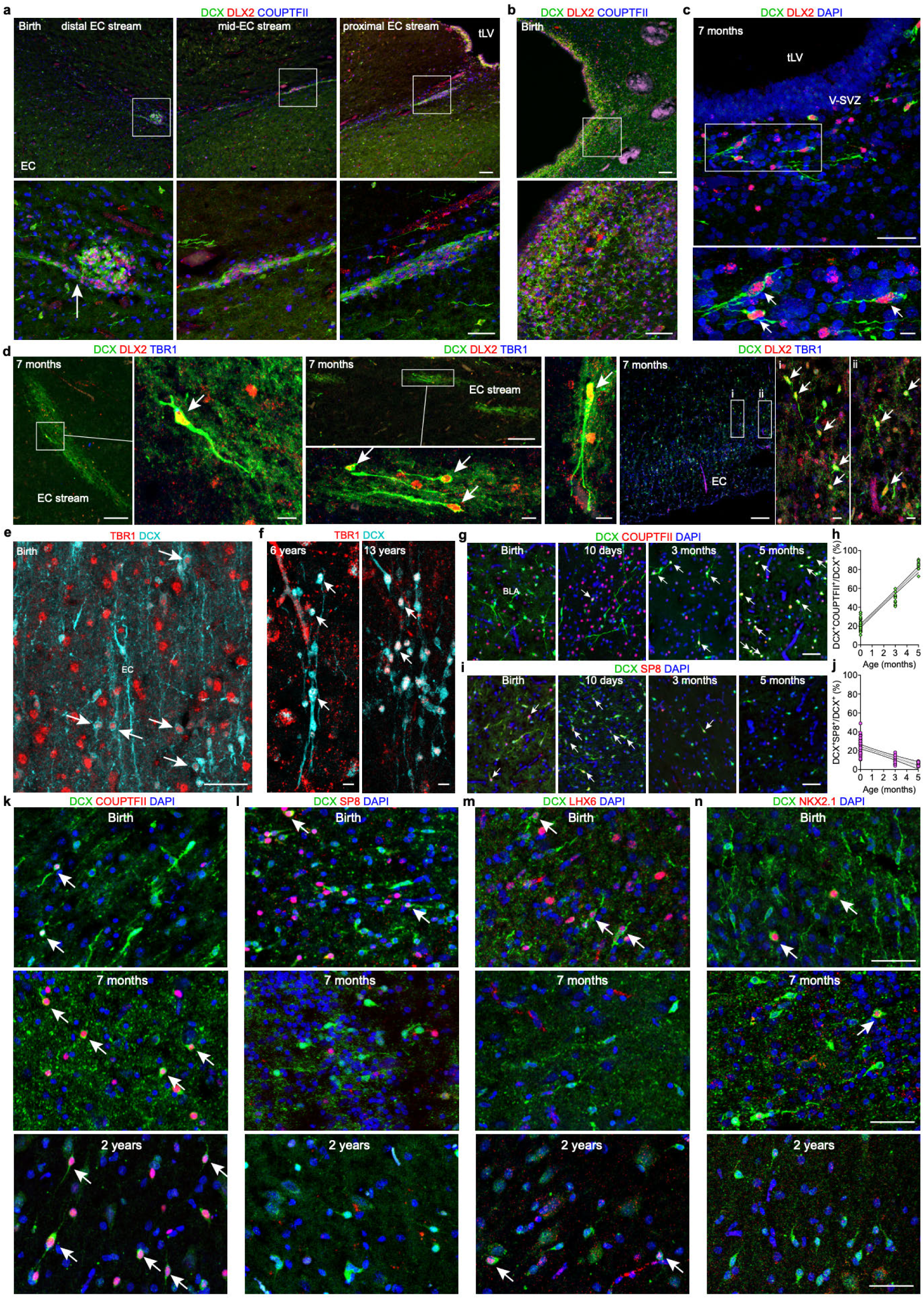
The EC stream supplies interneurons to the temporal lobe. **a**, DCX^+^ cells in the EC stream clusters at birth are DLX2^+^COUPTFII^+^ at distal, middle, and proximal distances from the tLV. Cells can be observed sending processes out of the dense cell clusters toward the EC (arrow). **b**, In the same section as (**a**), the lateral wall of the tLV contains large collections of DCX^+^DLX2^+^COUPTFII^+^ cells. **c**, At 7 months of age, a few individual DCX^+^DLX2^+^ cells are found ventrally to the tLV (arrows). **d**, At 7 months the EC stream and EC both contain DCX^+^DLX2^+^TBR1^−^ cells (arrows). **e**, In the EC at birth, DCX^+^ cells can be observed that co-express the cortical excitatory neuron transcription factor TBR1, however these cells typically had a more rounded morphology (arrows) and lacked migratory features. **f**, At 6 years and 13 years, individual DCX^+^TBR1^+^ neurons with simple morphology and small somas (arrows) persist in the EC primarily within layer 2. **g**, The neonatal basolateral amygdala also contains individual DCX^+^ cells at expressing COUPTFII with increasingly higher frequency by 5 months of age (arrows). **h**, Quantifications of the percentage of DCX^+^ cells co-expressing COUPTFII^+^ in the basolateral amygdala shows that this fraction rapidly predominates in the postnatal temporal lobe. (n= 4 cases at birth, 1 at 3 months and 1 at 5 months). **i**, The basolateral amygdala contains fewer individual DCX^+^ cells co-expressing SP8 at birth (arrows) and this fraction diminishes postnatally. **j**, Quantifications of the percentage of DCX^+^ cells co-expressing SP8 in the basolateral amygdala shows that this fraction declines between birth and 5 months. Quantifications are from adjacent sections to those in (**h**). **k–n**, Individual DCX^+^ cells in the EC at birth initially co-express (arrows) the transcription factors (**k**) COUPTFII, (**l**) LHX6, (**m**) SP8, (**n**) or NKX2.1. At the older ages of 7 months and 2 years, individual DCX^+^ cells in the EC predominantly express COUPTFII. Scale bars: 100 µm (**a** top, **b** top, **d** overviews), 50 µm (**a** bottom, **b** bottom, **c** top, **e, g, i, k–m**), 10 µm (**c** bottom, **d** all insets, **f**).

**Extended Data Fig. 8:**
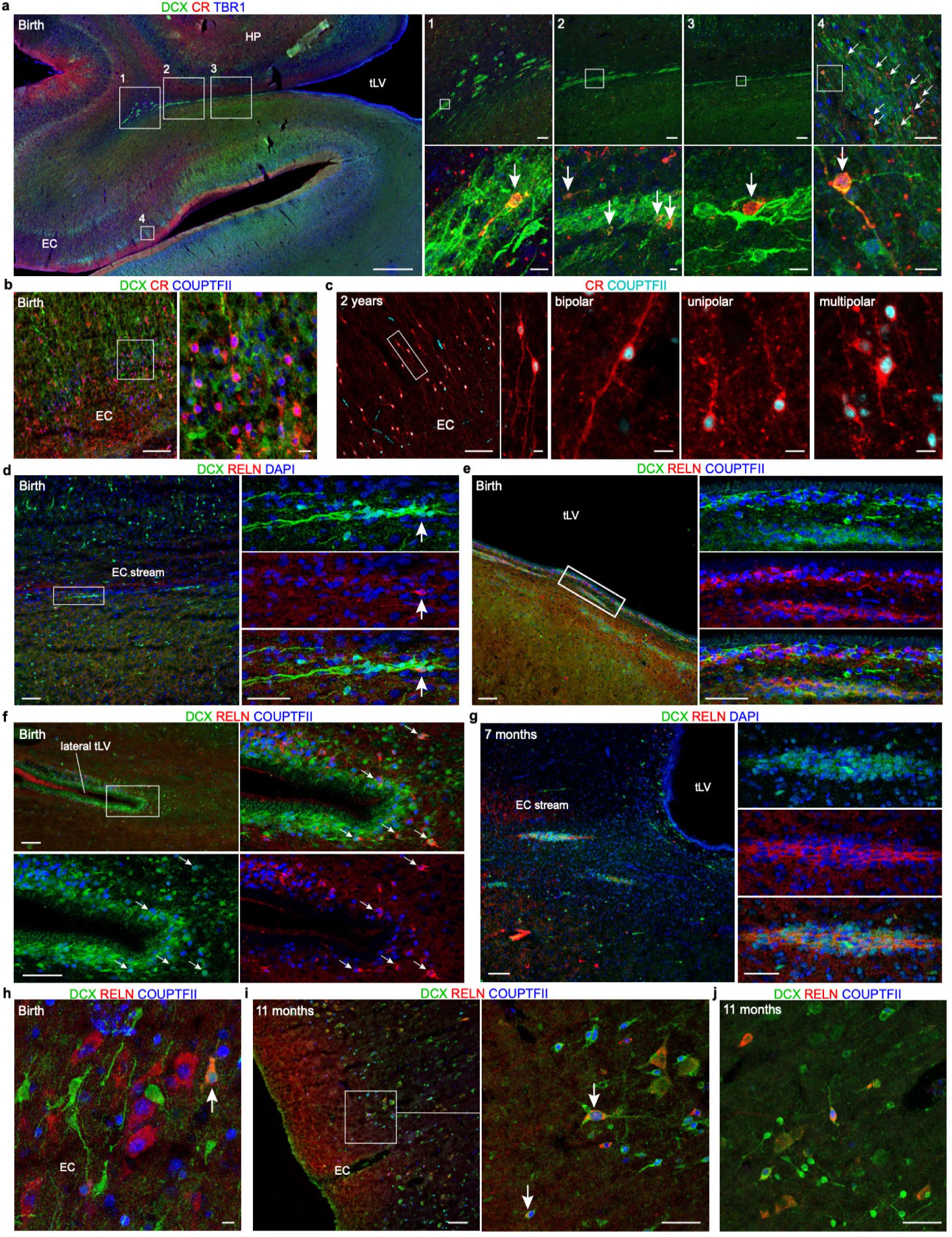
The EC stream supplies CR^+^ and RELN^+^ interneurons to the EC. **a**, Reproduction of the coronal section shown at lower magnification in **Fig. 4e**, with additional high magnification insets of DCX^+^CR^+^TBR1^−^ cells (arrows) in the EC stream clusters and EC (1-4). These cells are detectable in the EC stream at distal (1), middle (2), and proximal (3) levels relative to the medioventral tLV where the stream breaks away from the ventricle. Many additional DCX^+^CR^+^TBR1^−^ cells (arrows) are present in the EC (4) at birth, at the putative termini of the stream. **b**, At birth the EC contains DCX^+^CR^+^COUPTFII^+^ cells. **c**, At 2 years of age the vast majority of CR^+^ neurons in the EC co-express COUPTFII (inset). These cells display multiple morphologies including unipolar, bipolar, and multipolar. **d–f**, At birth only a few individual DCX^+^RELN^+^ neurons (arrow) are present in the EC stream (**d**), whereas the ventral wall of the tLV contains multiple chains of DCX^+^RELN^+^COUPTFII^+^ neurons (**e**), some of which are observed in the DCX^+^ chains wrapping around the lateral edge of the tLV (**f**). **g**, At 7 months of age, DCX^+^RELN^+^ cells are present in some of the dense clusters within the EC stream. **h**, At birth, the EC contains DCX^+^RELN^+^COUPTFII^+^ cells (arrow). **i**, At 11 months, similar DCX^+^RELN^+^COUPTFII^+^ cells are present in the EC, adjacent to DCX^+^COUPTFII^+^ neurons. **j**, Larger field of view of DCX^+^RELN^+^COUPTFII^+^ neuron in **Fig. 4n** at 11 months. Scale bars: 1 mm (**a** left), 100 µm (**a** 1–3 top, **b** left, **c** left, **d** left, **e** left, **f** top left, **g** left, **i** left), 50 µm (**a** 4 top, **d** right insets, **e** right insets, **f** bottom and right insets, **g** right insets, **i** right, **j**), 10 µm (**a** 1–4 bottom, **b** right, **c** right, **h**).

**Extended Data Fig. 9:**
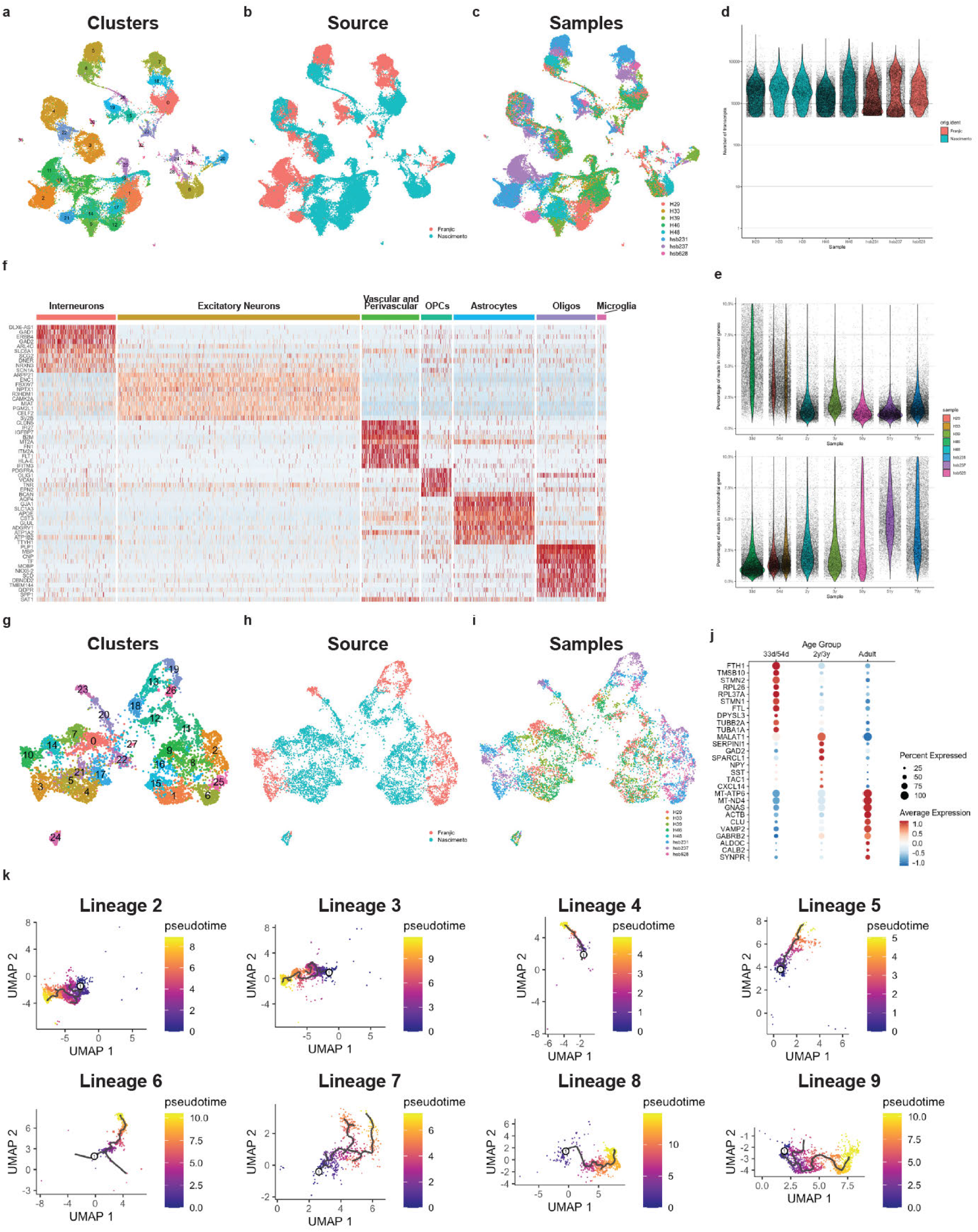
snRNA-seq dataset composition. **a-c**, UMAP plots of the integrated dataset, highlighting the assigned clusters (**a**), their dataset of origin (**b**) or sample of origin (**c**). **d**, Violin plot showing the number of reads/cell in each sample, colored by dataset of origin. **e**, Violin plots of the percentage of reads originating from ribosomal genes (upper panel) and mitochondrial genes (lower panel), colored by sample of origin and sorted by age. **f**, Heatmap of the top DEGs for each cell identity. **g-i**, UMAP plots of the reclustered interneurons, showing their assigned clusters (**g**), dataset of origin (**h**) and sample of origin (**i**). **j**, Dotplot of the top DEGs for inteneurons in each age group. **k**, UMAP plots showing the inferred lineage trajectories and estimated pseudotime for each interneuron lineage.

**Extended Data Fig. 10:**
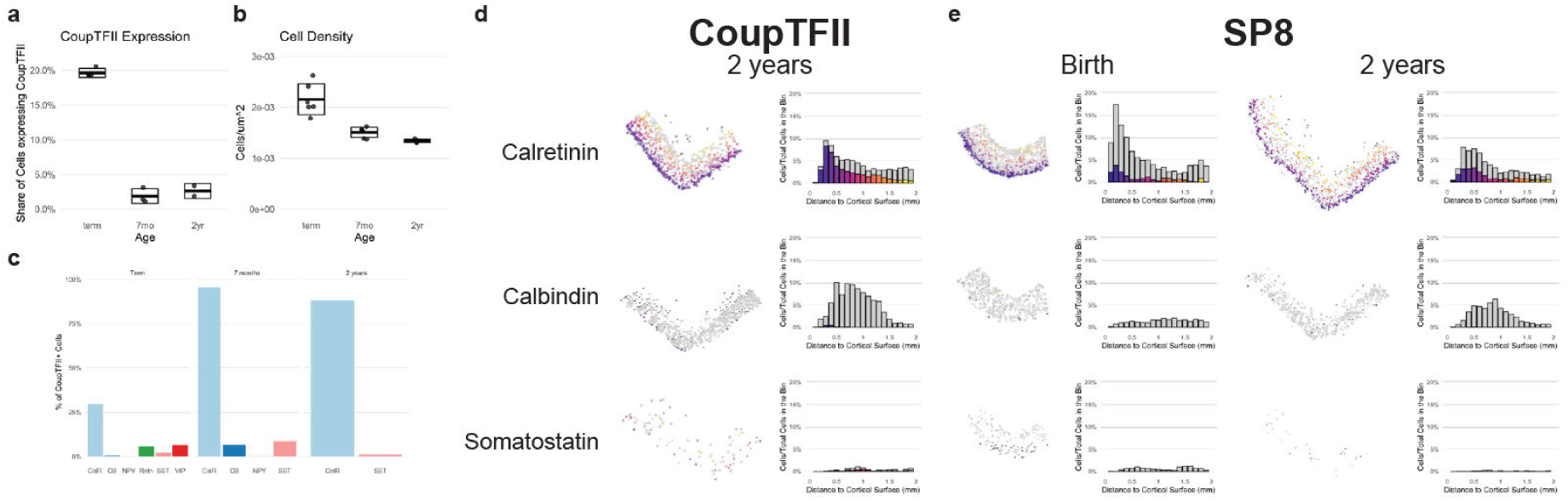
Extended data on CGE-derived interneurons in the EC. **a-b**, Percentage of cells expressing COUPTFII in the EC (**a**) and the overall cell density in the EC (**b**) at birth, 7 months and 2 years of age. **c**, Quantification of COUPTFII+ cells expressing different interneuron markers in the infant EC. **d-e**, Mapping and layer distribution of calretinin+ (upper row), calbindin+ (middle row) and somatostatin+ (lower panel) cells and its co-expression with COUPTFII+ at 2 years of age (**d**) and with SP8 at birth and 2 years of age (**e**).

